# A bacterial “shield and sword”: A previously uncharacterized two-component system protects uropathogenic *Escherichia coli* from host-derived oxidative insults and promotes hemolysin-mediated host cell pyroptosis

**DOI:** 10.1101/2021.04.19.440293

**Authors:** Hongwei Gu, Xuwang Cai, Xinyang Zhang, Jie Luo, Xiaoyang Zhang, Wentong Cai, Ganwu Li

**Author notes:** These authors contribute to this work equally (Hongwei Gu, Xuwang Cai).

## Abstract

Uropathogenic *Escherichia coli* (UPEC) deploys an array of virulence factors to successfully establish urinary tract infections. Coordinated expression of these various virulence factors is critical for UPEC’s overall fitness in the host. Two-component signaling systems (TCSs) are a major mechanism by which bacteria sense environmental cues and initiate adaptive responses. Here, we report a previously uncharacterized TCS encoded on a pathogenicity island in UPEC that directly activates the expression of a putative methionine sulfoxide reductase system (C3566/C3567) and a pore-forming α-hemolysin in response to host-derived hydrogen peroxide (H_2_O_2_) exposure. The TCS increases UPEC resistance to H_2_O_2_ *in vitro* and survival in macrophages in tissue culture via C3566/C3567. Additionally, the TCS mediates α-hemolysin-induced renal epithelial cell and macrophage death via a pyroptosis pathway. Taken together, our data suggest a paradigm in which this signal transduction system coordinates both bacterial pathogen defensive and offensive traits in the presence of host-derived signals.

## Introduction

Uropathogenic *Escherichia coli* (UPEC) cause most urinary tract infections (UTIs) worldwide. Cystitis and pyelonephritis are common clinical manifestations of UTIs, and in some severe cases, bacteremia occurs^1, 2^. Upon entry into the urethral opening, UPEC can ascend through the urethra and finally arrive at the bladder, where they can colonize and establish infection^3^. Immune cells, such as macrophages and neutrophils, function as the first-line barriers to UPEC infection, usually by engulfing and killing the bacteria with reactive oxygen (ROS, e.g., H_2_O_2_), chlorine (RCS, e.g., HOCl) and nitrogen species (RNS, e.g., NO)^4–6^. However, pathogenic bacteria can resist hostile immune killing, which can be deemed defensive activity. Defense systems in UPEC include the KatE and KatG catalases that decompose H_2_O_2_^7^, and RpoS^8^ regulator that elicit global response to withstand oxidative stress. As another strategy for survival, many bacterial pathogens encode toxins that can damage host cells in various ways^9, 10^, which can be deemed offensive activity. Thus, an effective and efficient coordination of defense and offense during infection is conceivably an important fitness factor for pathogens.

UPEC form a very diverse group of bacteria possessing a large repertoire of virulence factors^11^, including hemolysin. Hemolysin is a prototypic alpha pore-forming toxin that is carried by approximately half of UPEC clinical isolates, with an even greater percentage in pyelonephritis isolates. Clinically, hemolysin carriage is associated with severe symptoms, especially kidney damage^12, 13^. Hemolysin has been proposed to function differentially over a gradient of concentrations. At high concentrations, it can form pores in membranes of various host cell types, including red blood cells (RBCs), epithelial cells and leukocytes, resulting in cell lysis^14–16^. However, at low (sublytic) concentrations, hemolysin can either alter cell functions or induce cell death. For example, in UPEC str. UTI89, sublytic doses of hemolysin can activate host proteases, especially mesotrypsin, resulting in degradation of paxillin that aids in host cell membrane damage and further detachment of the cell^17^. One study reported that hemolysin from UTI89 activates a caspase-1/caspase-4-dependent death pathway, most likely pyroptosis, in human bladder epithelial cells^18^. In addition, hemolysin from the blood isolate CP9 from a patient with sepsis has been found to mediate apoptosis and necrosis of human neutrophils^15^. Despite the well-documented toxic functions of hemolysin, in-depth studies on its regulation are very limited. Reported regulators include CpxR^18^, FNR^19^, RfaH^20^ and BarA/UvrY^21^, and several other factors have been identified in genetic screens, including Cof, which is a phosphatase^14^; LPS biosynthesis gene products; and DnaKJ chaperones^22^. Importantly, the settings in which these factors contribute are insufficiently studied.

Two-component signaling systems (TCSs), which are composed of a membrane-bound histidine kinase (HK) sensor and a cytoplasmic response regulator (RR), have been implicated in regulating bacterial responses to a variety of signals and stimuli, such as nutrients and small-molecule signals. Recognition of physical or chemical signals by the HK sensor domain typically triggers modulation of HK autophosphorylation activity. The phosphoryl group is then transferred to the RR, which is usually a DNA-binding protein that functions by altering gene expression^23, 24^. Our group have previously revealed that UPEC-associated TCS KguS/R, which respond to the presence of α-ketoglutarate (KG), a particularly abundant metabolite within renal proximal tubule cells, stimulate the expression of KG utilization genes^25^. As UPEC reside in distinct host environments compared to enteric *E. coli*^26^, it seems plausible that UPEC carry a particular set of TCSs for environmental adaptation and virulence.

In this study, our group identified a previously uncharacterized genomic island-encoded TCS, *c3564*/*c3565*, that is highly associated with UPEC. In response to H_2_O_2_ exposure *in vitro* and within macrophages, C3564/C3565 directly activates the expression of *c3566*-*c3567*, which encodes a putative methionine sulfoxide (MetSO) reductase system (Msr) that protect the bacteria from oxidative insults, increasing survival in oxidative environments and in host cells. In addition, C3564/C3565 promoted hemolysin-mediated host cell pyroptosis in the presence of H_2_O_2_. Mechanistically, C3564/C3565 coordinates oxidative resistance and toxin production by cotranscribing the *c3566*-*c3567* and *hlyCABD* operons. Overall, our study provides a paradigm in which a regulatory system links together defensive and offensive activities in a bacterial pathogen.

## Results

### C3564 and C3565 constitute a cognate TCS

In searching for previously uncharacterized TCSs in pathogenic *E. coli*, we found that *c3564* and *c3565*, which are located in the *pheV* genomic island in UPEC CFT073, are predicted to encode a TCS. C3564 is predicted to be an HK that contains a periplasmic domain flanked by a transmembrane helix on each side, a dimerization/phosphoacceptor (HisKA) domain containing a conserved autophosphorylation site (histidine, His278), and an ATPase domain (Fig. 1a). C3565 is predicted to be an OmpR subfamily RR possessing a CheY-like receiver domain containing an aspartate residue (Asp55) presumed to accept a phosphoryl group from an HK, as well as a winged helix-turn-helix (HTH) DNA-binding domain (Fig. 1a).

**Figure 1.**
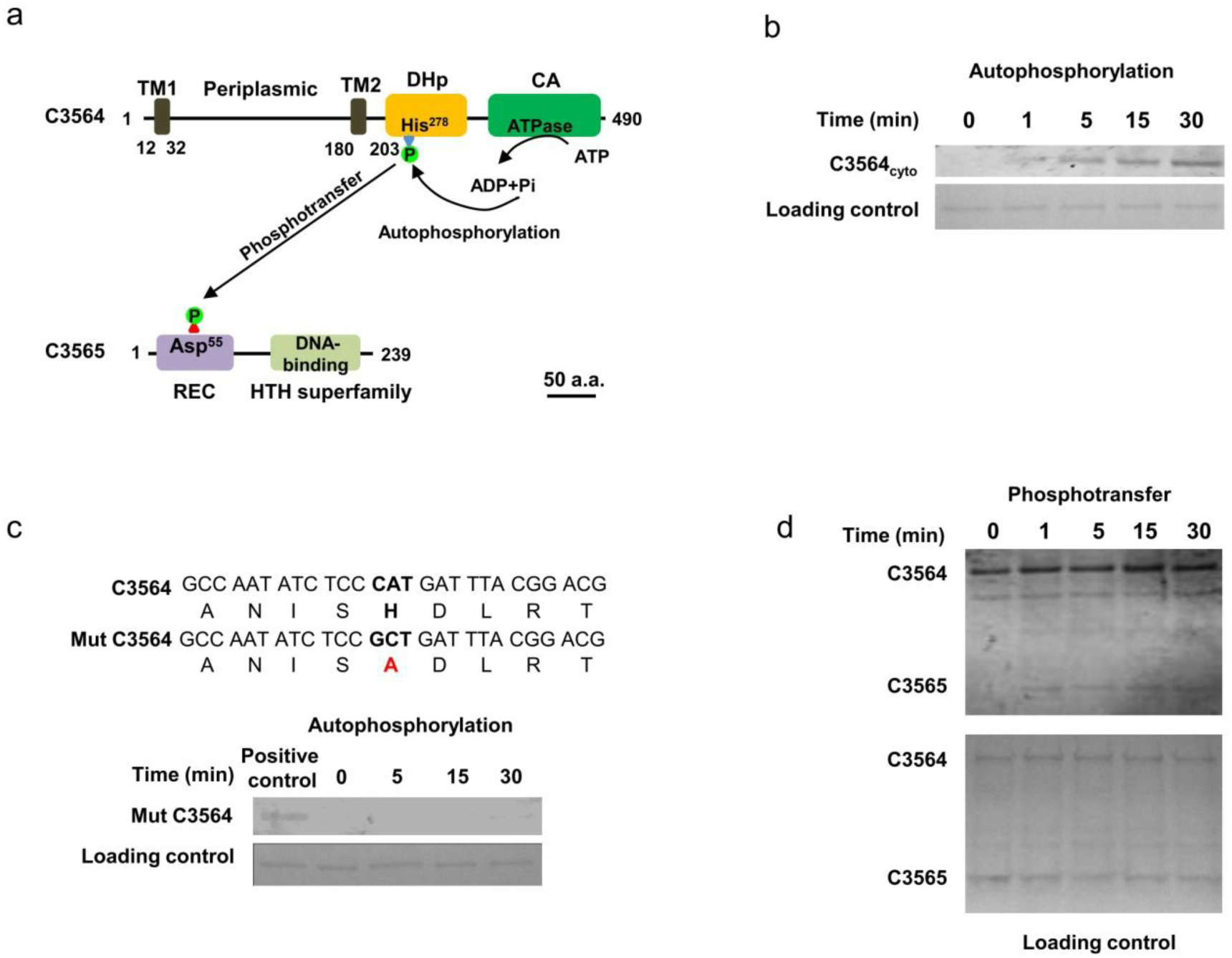
Identification of a two-component signaling system, C3564/C3565, in UPEC CFT073. **a**, Domain architectures of C3564 and C3565. The numbers indicate the amino acid positions. The domains were predicted with the InterPro database. Abbreviations: TM1 and TM2, two transmembrane helices; DHp, dimerization and histidine phosphorylation domain containing His278 for autophosphorylation; CA, catalytic ATPase domain; REC, receiver domain containing the putative aspartate residue for receiving the phosphoryl group; HTH, helix-turn-helix domain for DNA binding. The scale bar indicates 50 amino acids (a.a.). **b**, *In vitro* analysis of C3564 autophosphorylation. The purified cytoplasmic portion of C3564 was incubated with ATP for various amounts of time, and phosphorylated proteins were detected using the pIMAGO-biotin phosphoprotein detection method. **c**, Mutation of His278 abolished C3564 self-phosphorylation. Mut C3564 is a variant of C3564 carrying a His278Ala mutation, and the positive control is the wild-type C3564 protein. **d**, *In vitro* transphosphorylation of C3565 by phosphorylated C3564. The autophosphorylated form of C3564 and purified C3565 protein were mixed at equimolar concentrations and then incubated at 37°C for the indicated amounts of time. The reaction mixture was directly subjected to SDS-PAGE and detected by the pIMAGO-biotin phosphoprotein detection method. These experiments were repeated at least twice, and representative images are shown.

To confirm whether C3564 and C3565 constitute a cognate TCS, autophosphorylation and transphosphorylation assays were performed with purified recombinant proteins. C3564 was expressed as a cytoplasmic portion (C3564_cyto_)-GST fusion protein, and C3565 as a His_6_-C3565 fusion protein. Autophosphorylation occurred rapidly (at 1 min) and increased as the incubation was prolonged (Fig. 1b). In contrast, when a mutant form of C3564 in which the conserved His278 was changed to alanine (His278Ala) was tested, no autophosphorylation was detected within the 30 min time frame of the assay (Fig. 1c), suggesting that His278 is involved in accepting the phosphoryl group. A transphosphorylation assay was performed using ATP-saturated C3564 coincubated with C3565. We observed markedly increased levels of C3565 phosphorylation within 1 min compared to the levels for the control protein at 0 min (Fig. 1d), indicating that C3565 can be transphosphorylated by C3564. Together, these results suggest that C3564 and C3565 constitute a cognate TCS.

### *c3564* and *c3565* are highly associated with UPEC and promote host cell death during infection in cell culture models

To examine the prevalence of *c3564*/*c3565* in pathogenic *E. coli* strains, we first performed a BlastN search analysis, and the results demonstrated that *c3564*/*c3565* orthologs can be found in most UPEC genomes available in the NCBI database but not in other pathotypes, such as avian pathogenic *E. coli* (APEC), EHEC, and enterotoxigenic *E. coli* (ETEC). Duplex PCR detection of *c3564* and *c3565* in a large collection of *E*. *coli* clinical isolates showed that these two genes are highly associated with the uropathogenic pathotype (Fig. 2a), implying that *c3564*/*c3565* might be important for UPEC pathogenicity. We then constructed two single-gene deletion mutants of CFT073, Δ*c3564* and Δ*c3565*, and used them to examine whether *c3564* and *c3565* play a role in the interaction between UPEC and host cells. Pronounced cytopathic effects, including cell rounding and detachment, were observed in kidney epithelial cells (A498) after 3 h of infection with wild-type CFT073 (Fig. 2b). In contrast, deletion of either *c3564* or *c3565* nearly eliminated the cytopathic effects caused by CFT073, suggesting that *c3564* and *c3565* are involved in damaging host cells. Next, we quantified cell death by using an LDH release assay. Infection with wild-type CFT073 led to substantial A498 cell death, whereas infection with the Δ*c3564* and Δ*c3565* mutants resulted in significantly less cell death (Fig. 2c). Introduction of a plasmid-borne *c3564* or *c3565* locus into the corresponding mutant restored the cytopathic effects caused by CFT073 (Fig. 2c). Additionally, in two other cell types, the human macrophage-like cell line THP-1 and the mouse macrophage cell line J774A.1, deletion of *c3564* or *c3565* significantly reduced UPEC-induced host cell death (Fig. 2d and e). Altogether, these data demonstrate that *c3564* and *c3565* promote host cell death during UPEC infection *ex vivo*.

**Figure 2.**
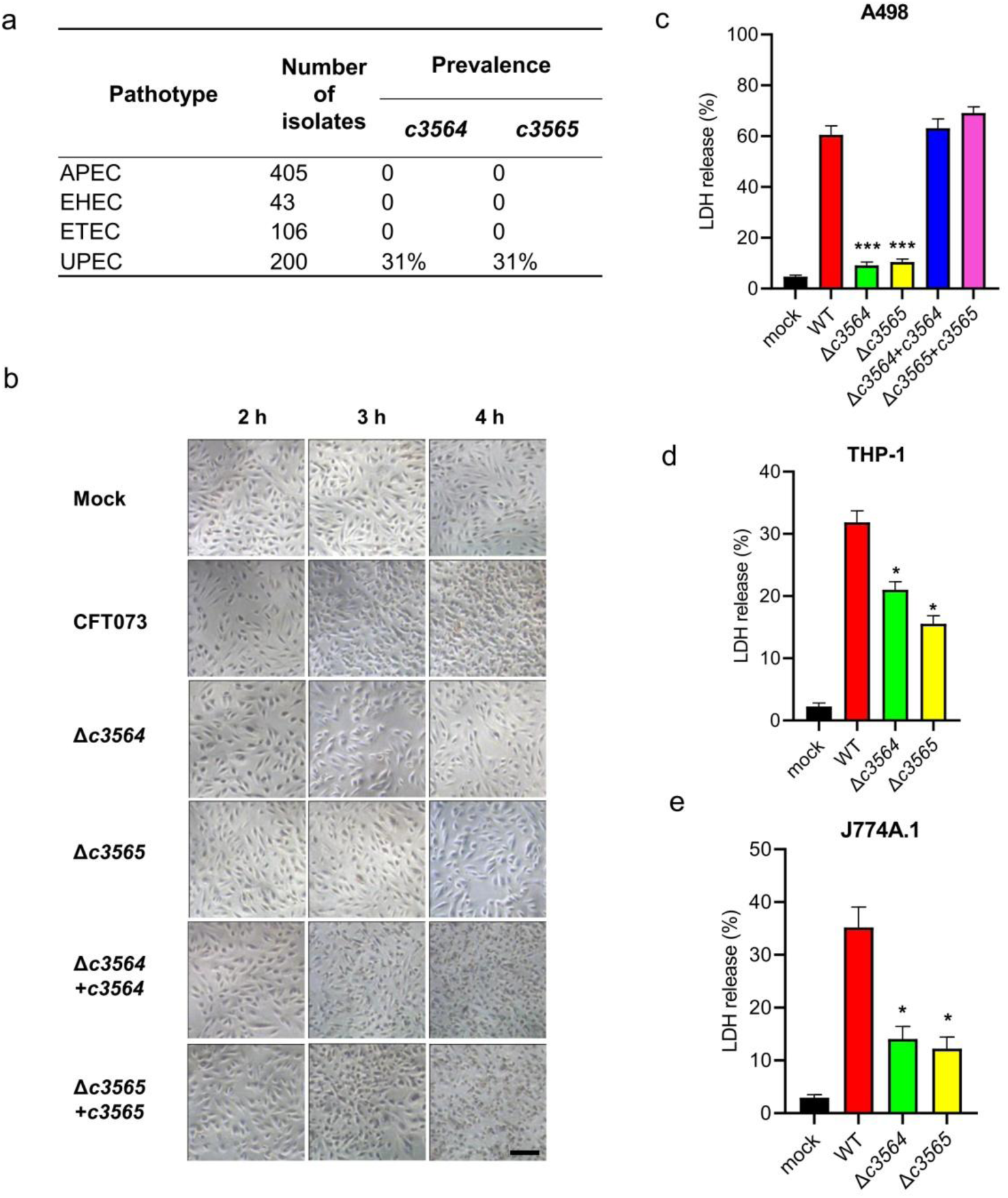
*c3564* and *c3565* promote host cell death during infection in cell culture models. **a,** Prevalence of *c3564* and *c3565* in various *E. coli* pathotypes. A duplex PCR method was used to detect the presence of the *c3564* and *c3565* genes in a laboratory collection of *E. coli* isolates. APEC, avian pathogenic *E. coli*; EHEC, enterohemorrhagic *E. coli*; ETEC, enterotoxigenic *E. coli*; UPEC, uropathogenic *E. coli*. **b**, *c3564* and *c3565* affected the morphology of kidney epithelial cells. A498 kidney epithelial cells were infected with CFT073 and its derivatives for the indicated amounts of time and then subjected to phase-contrast microscopy (magnification, 20×). Scale bar, 50 μm. **c,d and e**, Cytotoxicity assays on different cell types. A498 kidney epithelial cells (c), THP-1 human macrophages (d), and J774A.1 murine macrophage cells (e) were infected with various bacterial strains at a multiplicity of infection (MOI) of ∼10 for 2.5 h, and the cell culture supernatants were then subjected to LDH release measurement. Cytotoxicity (%) was determined by comparing the LDH in culture supernatants to the total cellular LDH (the amount of LDH released upon cell lysis with 0.1% Triton X-100) according to the formula [(experimental − target spontaneous)/(target maximum − target spontaneous)] × 100. The data are the mean ± SD from three independent experiments. *, *P <* 0.05; ***, *P <* 0.001 by one-way ANOVA followed by Dunnett’s multiple comparisons test against wild-type CFT073.

### C3564 and C3565 mediate induction of host cell pyroptosis through controlling hemolysin production

To elucidate the mechanism by which C3564 and C3565 promote host cell death, RNA-seq analysis was performed to identify genes regulated by C3564 and C3565 during bacterium-cell interactions. Wild-type CFT073 or its mutants associated with A498 cells were collected, and bacterial RNA extraction and transcriptome sequencing were then performed. A 2-fold change was selected as the cutoff. Table 1 lists some of the genes that were most downregulated due to deletion of *c3564* or *c3565*. Notably, the *hlyCABD* genes, responsible for the production and translocation of hemolysin, were downregulated by ∼4-fold or more in the Δ*c3564* and Δ*c3565* mutants. Furthermore, immunoblotting and blood agar plate hemolysis assays confirmed that deletion of either *c3564* or *c3565* impaired hemolysin production and subsequent hemolysis, and these phenotypic changes can be properly complemented (Fig. 3a). These results indicate that C3564 and C3565 positively regulate hemolysin production.

**Figure 3.**
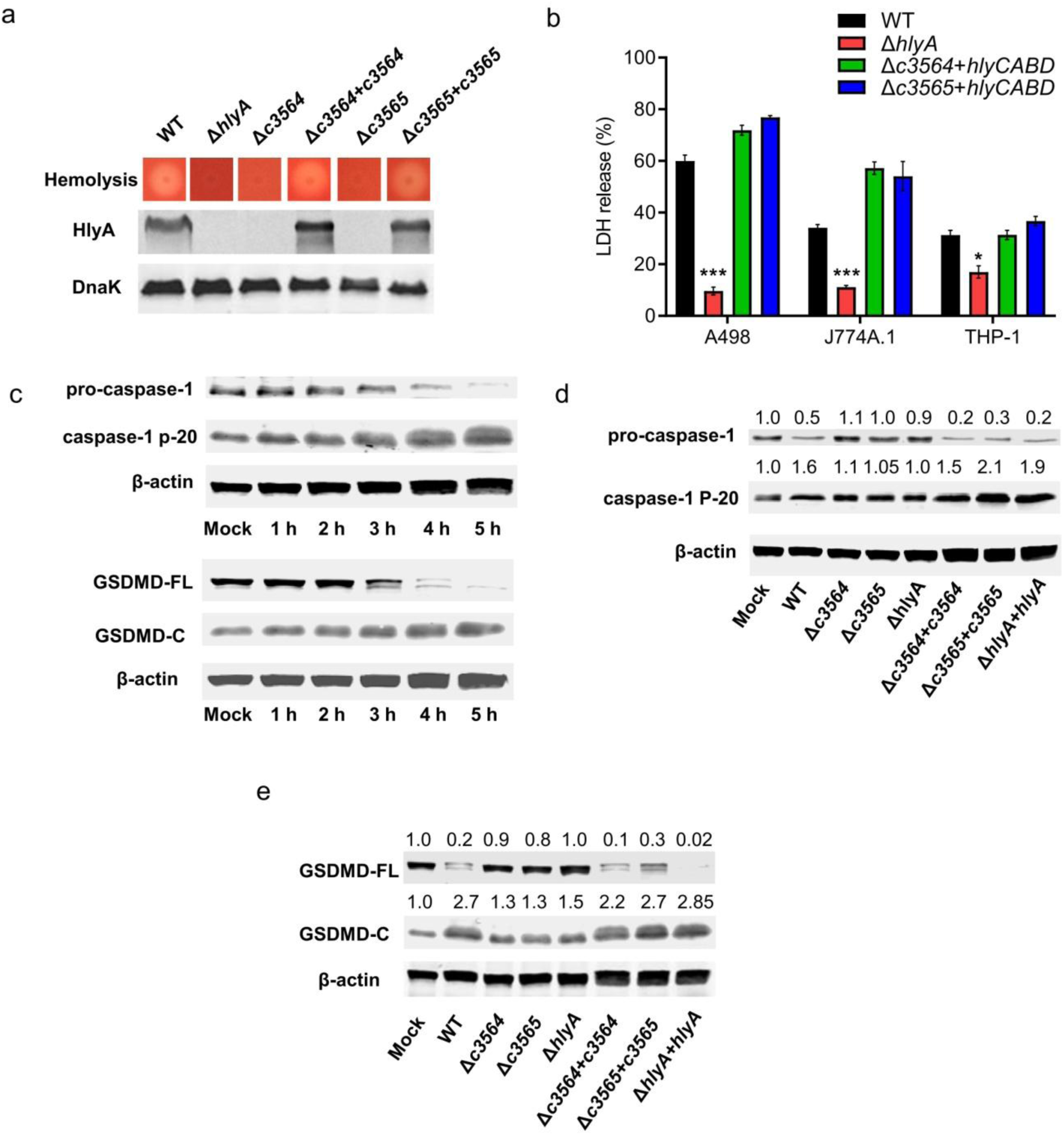
*c3564* and *c3565* promote hemolysin-mediated host cell pyroptosis. **a**, *c3564* and *c3565* regulate hemolysin production and hemolysis on sheep blood agar. The dark dot in the center of each image represents a bacterial colony, and the halo around the dark dot represents hemolysis. A wider halo indicates stronger hemolysis. **b**, *c3564-* and *c3565*-mediated hemolysin-induced host cell death in different host cell types. *, *P <* 0.05; ***, *P <* 0.001 by one-way ANOVA followed by Dunnett’s multiple comparisons test against wild-type CFT073. **c**, UPEC CFT073 induces the processing of pro-caspase-1 and GSDMD. **d and e,** *c3564* and *c3565* promote hemolysin-mediated processing of pro-caspase-1 (d) and GSDMD (e) during infection of A498 cells. The numbers above the blots indicate the relative expression levels, with that of the mock group set as 1.0, based on densitometry with ImageJ. All blots are representative of ≥2 independent experiments.

**Table 1.**
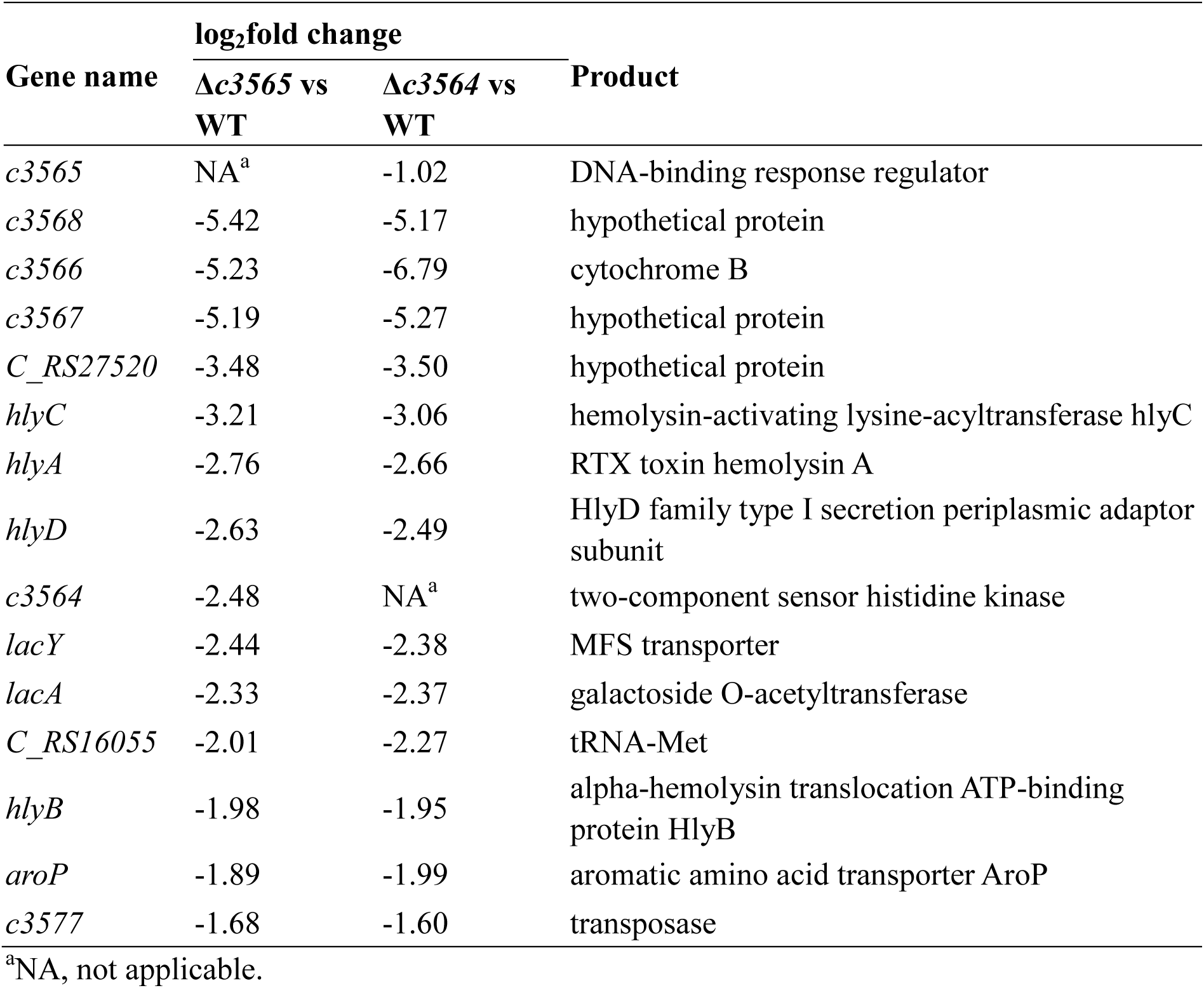
Select genes that were differentially expressed in the Δ*c3564* and Δ*c3565* mutants compared to the wild type.

| Gene name | log<sub>2</sub>fold change |  | Product |
| --- | --- | --- | --- |
| | $\Delta c3565$ vs WT | $\Delta c3564$ vs WT | |
| *c3565* | NA<sup>a</sup> | -1.02 | DNA-binding response regulator |
| *c3568* | -5.42 | -5.17 | hypothetical protein |
| *c3566* | -5.23 | -6.79 | cytochrome B |
| *c3567* | -5.19 | -5.27 | hypothetical protein |
| *C_RS27520* | -3.48 | -3.50 | hypothetical protein |
| *hlyC* | -3.21 | -3.06 | hemolysin-activating lysine-acyltransferase hlyC |
| *hlyA* | -2.76 | -2.66 | RTX toxin hemolysin A |
| *hlyD* | -2.63 | -2.49 | HlyD family type I secretion periplasmic adaptor subunit |
| *c3564* | -2.48 | NA<sup>a</sup> | two-component sensor histidine kinase |
| *lacY* | -2.44 | -2.38 | MFS transporter |
| *lacA* | -2.33 | -2.37 | galactoside O-acetyltransferase |
| *C_RS16055* | -2.01 | -2.27 | tRNA-Met |
| *hlyB* | -1.98 | -1.95 | alpha-hemolysin translocation ATP-binding protein HlyB |
| *aroP* | -1.89 | -1.99 | aromatic amino acid transporter AroP |
| *c3577* | -1.68 | -1.60 | transposase |

<sup>a</sup>NA, not applicable.

LDH release assays further showed that an *hlyA* mutant produced limited cell death and that introduction of a plasmid carrying *hlyCABD* into Δ*c3564* or Δ*c3565* greatly restored the cytopathic effects (Fig. 3b), confirming that the TCS mediated induction of host cell death through controlling bacterial hemolysin production. To gain insights into the cell death pathways triggered when *c3564* and *c3565* are present, we infected A498 cells with wild-type CFT073 and its derivative strains, and then used flow cytometry to detect annexin V-FITC/PI staining of the infected cell. Our results showed that within 2 hpi the majority of cells became double-positive stained (Fig. S1), which is different from necrosis and apoptosis status, suggesting that the infected cells have undergone death pathways like pyroptosis other than apoptosis^27^. And deletion of *c3564*, *c3565*, or *hlyA* reduced annexin V/PI staining, suggesting that these genes are involved in this cell death phenotype (Fig. S1). Further, we found that in A498 kidney epithelial cells infected with wild-type CFT073, pro-caspase-1 was gradually processed into p20, and the production of p20 was increased significantly at 5 h post infection (hpi); in addition, full-length gasdermin D (GSDMD) was cleaved into GSDMD-N and GSDMD-C (Fig. 3c). As GSDMD is the effector molecule that determines cell pyroptosis^42^, these results indicate the onset of pyroptosis in A498 cells infected with the wild type. In contrast, deletion of *hlyA* in CFT073 significantly reduced the processing of pro-caspase-1 into caspase-1 (Fig. 3d) and the cleavage of GSDMD (Fig. 3e), indicating weakened pyroptosis induction by the mutant compared to that induced by the wild type. Thus, *hlyA* is important for CFT073-induced pyroptosis in kidney epithelial cells. When *hlyA* was present, deletion of either *c3564* or *c3565* reduced the occurrence of pyroptosis, and these phenotypic changes due to the gene deletions were able to be properly complemented (Fig. 3d and e). Furthermore, we also analyzed the roles of HlyA and C3564/C3565 in inducing pyroptosis in other host cell types, including THP-1 and J774A.1 macrophages, and the results showed that Δ*c3564*/*c3565* and Δ*hlyA* mutants presented diminished cleavage of full-length GSDMD, but introduction of *hlyCABD* genes into Δ*c3564*/*c3565* mutant markedly increased cleavage of full-length GSDMD (Fig. S2). Together, these results suggest that C3564/C3565 mediates induction of host cell pyroptosis through regulating bacterial hemolysin production.

### C3564 and C3565 control hemolysin expression by directly regulating *c3566*-*c3568*

Then, we began to explore the mechanism underlying the regulation of hemolysin by C3564/C3565. Among the differentially expressed genes in the Δ*c3564*/wild type and Δ*c3565*/wild type comparisons, *c3566*, *c3567*, and *c3568*, which are immediately downstream of *c3564*/*c3565*, were the genes most downregulated upon knockout (Table 1). Quantitative real-time PCR (qPCR) confirmed that C3564 and C3565 positively regulate the expression of *c3566*, *c3567*, and *c3568* (Fig. 4a). Reverse transcription PCR (RT-PCR) results showed that *c3566*, *c3567*, and *c3568* are cotranscribed and therefore form an operon (Fig. 4b). The transcription start site (TSS) of the *c3566*-*c3568* operon was determined by 5’-RACE PCR, and the ribosomal binding site (RBS) as well as the −10 and −35 boxes were deduced using BProm program (http://linux1.softberry.com) (Fig. 4b). In an electrophoretic mobility shift assay (EMSA) with a 200 nt DNA fragment containing the promoter region as a probe, we found that the C3565 protein directly binds to the promoter region (Fig. 4c). Indeed, a potential RR-binding region containing inverted repeats (TCACA-N_15_-TGTGA) was identified immediately upstream of the −35 box (Fig. 4b). Altogether, these data indicate that C3565 directly regulates the *c3566*-*c3568* operon.

**Figure 4.**
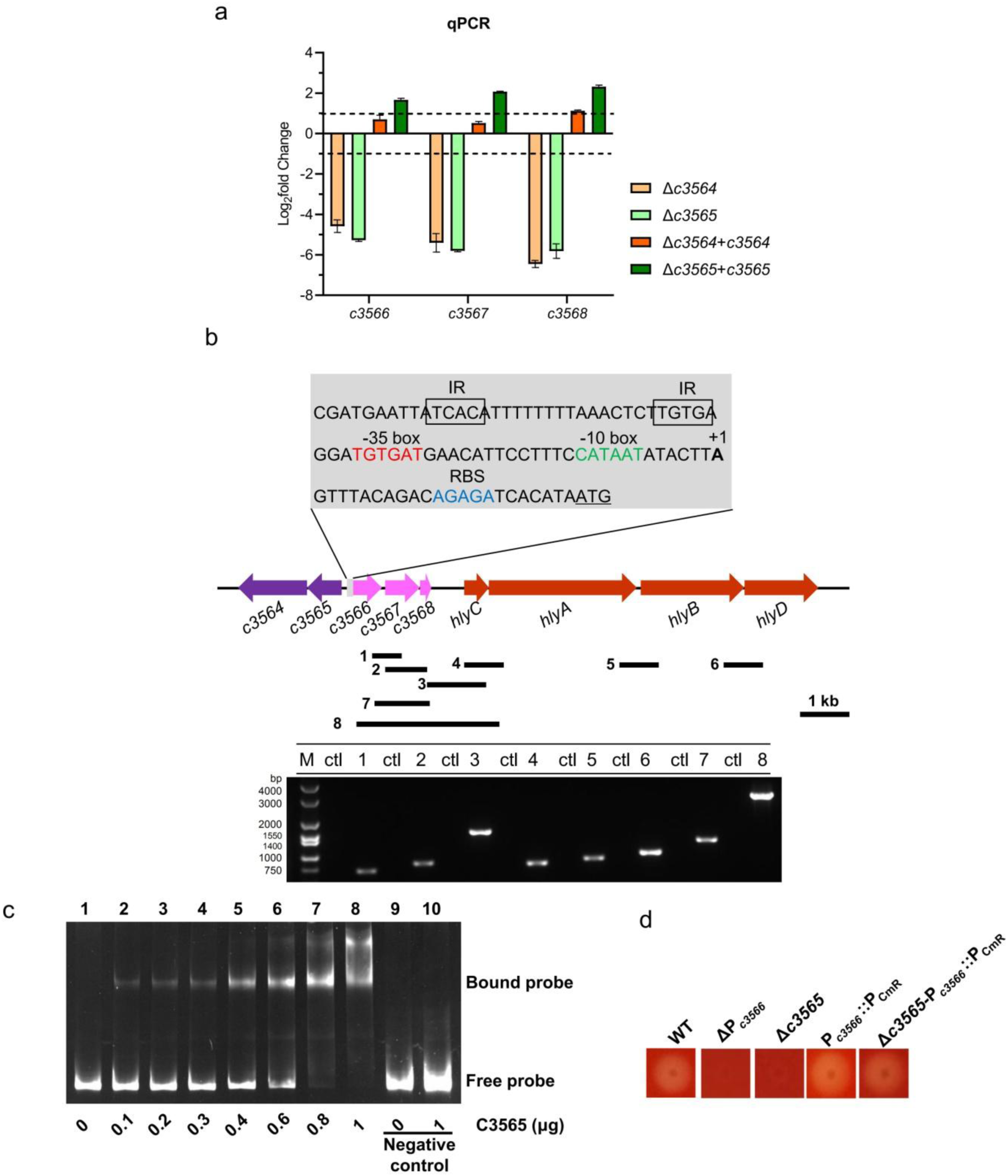
*c3566*-*c3568* transcription and *hlyCABD* transcription are linked and controlled by *c3564* and *c3565*. **a**, qPCR analysis indicates that *c3564* and *c3565* positively regulate the expression of *c3566*-*c3568* expression. The dashed lines represent the fold change cutoff of 2. A fold change ≥2 was considered significant. **b**, RT-PCR analysis indicates that the *c3566*-*c3568* operon and *hlyCABD* operon are co-transcribed. The black bars under the gene arrows denote the PCR amplicons generated during testing of cotranscription, and the number in front of each bar corresponds to the lane in the agarose gel. Ctl indicates control RNA without reverse transcription. 5’-RACE PCR was used to identify the TSS (+1) of the *c3566*-*c3568* operon, and the −10 and −35 motifs were deduced afterward. Gray box, promoter region of *c3566*; underlined ATG, start codon; blue letters, ribosomal binding site; green letters, −10 box; red letters, −35 box; boxed letters, inverted repeats (IRs) likely bound by the C3565 protein. The genes are drawn to scale. **c**, EMSA analysis of C3565 protein binding to the *c3566* promoter region. A fragment of CmR coding sequence was used as a negative control probe. **d**, Hemolysis is regulated by the *c3566* promoter (P*_c3566_*). Strain names represent: WT, wild-type CFT073; ΔP*_c3566_*, *c3566* promoter deletion mutant; Δ*c3565*, *c3565* deletion mutant; P*_c3566_*::P_CmR_, a mutant in which *c3566* promoter is replaced by P_CmR_; Δ*c3565*-P*_c3566_*::P_CmR_, a mutant in which *c3565* is deleted and *c3566* promoter is replaced by P_CmR_.

We then investigated how C3565 regulates the two adjacent operons, *c3566*-*c3568* and *hlyCABD*. An EMSA demonstrated that the C3565 protein does not associate with a region upstream of the *hlyA* coding sequence (−300 to +1 relative to the *hlyA* start codon) (data not shown), suggesting a lack of direct regulation. We therefore hypothesized that C3565 may regulate *hlyCABD* through other *cis-* or *trans*-elements. A series of deletion mutants were generated, including mutants with individual and combined deletions of *c3566*, *c3567*, and *c3568* and various deletions of the intergenic region between *c3568* and *hlyC*, but these genetic modifications did not affect hemolysis (Fig. S3). However, when the promoter P*_c3566_* (from the −35 box to the TSS) was deleted, hemolysis was lost. Insertion of a constitutively expressed P_CmR_ promoter in front of the *c3566* coding sequence restored hemolysis, and the restoration was independent of *c3565* (Fig. 4d). These results imply that *c3566*-*c3568* and *hlyCABD* can be cotranscribed. Indeed, RT-PCR demonstrated that *c3566*-*c3568*-*hlyCABD* can be transcribed as a polycistronic mRNA (Fig. 4b). Collectively, these results indicate that C3564 and C3565 control hemolysin expression by directly regulating *c3566*-*c3568-hlyCABD*.

### C3564/C3565 contributes to UPEC resistance to H_2_O_2_ *in vitro* as well as to survival within macrophages through C3566/C3567

Analysis of their amino acid sequences showed that C3567 and C3566 are 27.6% and 23.1% identical to *E. coli* MG1655 MsrP and MsrQ, respectively, which encode an Msr system. The periplasmic MsrP catalyzes the reduction of MetSO through its molybdenum-molybdopterin cofactor, while the membrane-bound MsrQ mediates electron supplies for the MetSO reduction likely using the quinone pool of the respiratory chain^32^. Methionine residues in proteins are particularly sensitive to oxidative stressors that can convert methionine into MetSO, which renders proteins dysfunctional^28, 29^. Msr system repairs methionine residues damaged by ROS and RCS, including H_2_O_2_. C3568 encodes a small methionine-rich peptide, and a homolog of C3568 in *Azospira*, MrpX, serves as an oxidative sacrificial sink protein that scavenges HOCl^31, 32^. Thus, we hypothesized that the TCS C3564/C3565 contributes to UPEC resistance to H_2_O_2_ through regulating *c3566*-*c3568* expression. In the presence of tert-butyl hydroperoxide (Ter-Bo), compared to the wild-type strain, separate *c3564*, *c3565*, *c3566* and *c3567* deletion mutants exhibited significantly reduced growth (Fig. 5a). In contrast, without Ter-Bo, no growth differences were observed between the wild-type and mutant strains. Complementation of *c3566* and *c3567 in trans* restored growth in the presence of Ter-Bo, and constitutive expression of *c3566*-*c3567* in the *c3564* or *c3565* mutant strains restored the growth to the wild-type level (Fig. 5a). These results indicate that the TCS C3564/C3565 promotes resistance to oxidative stress (Ter-Bo exposure) through regulating *c3566* and *c3567* expression.

**Figure 5.**
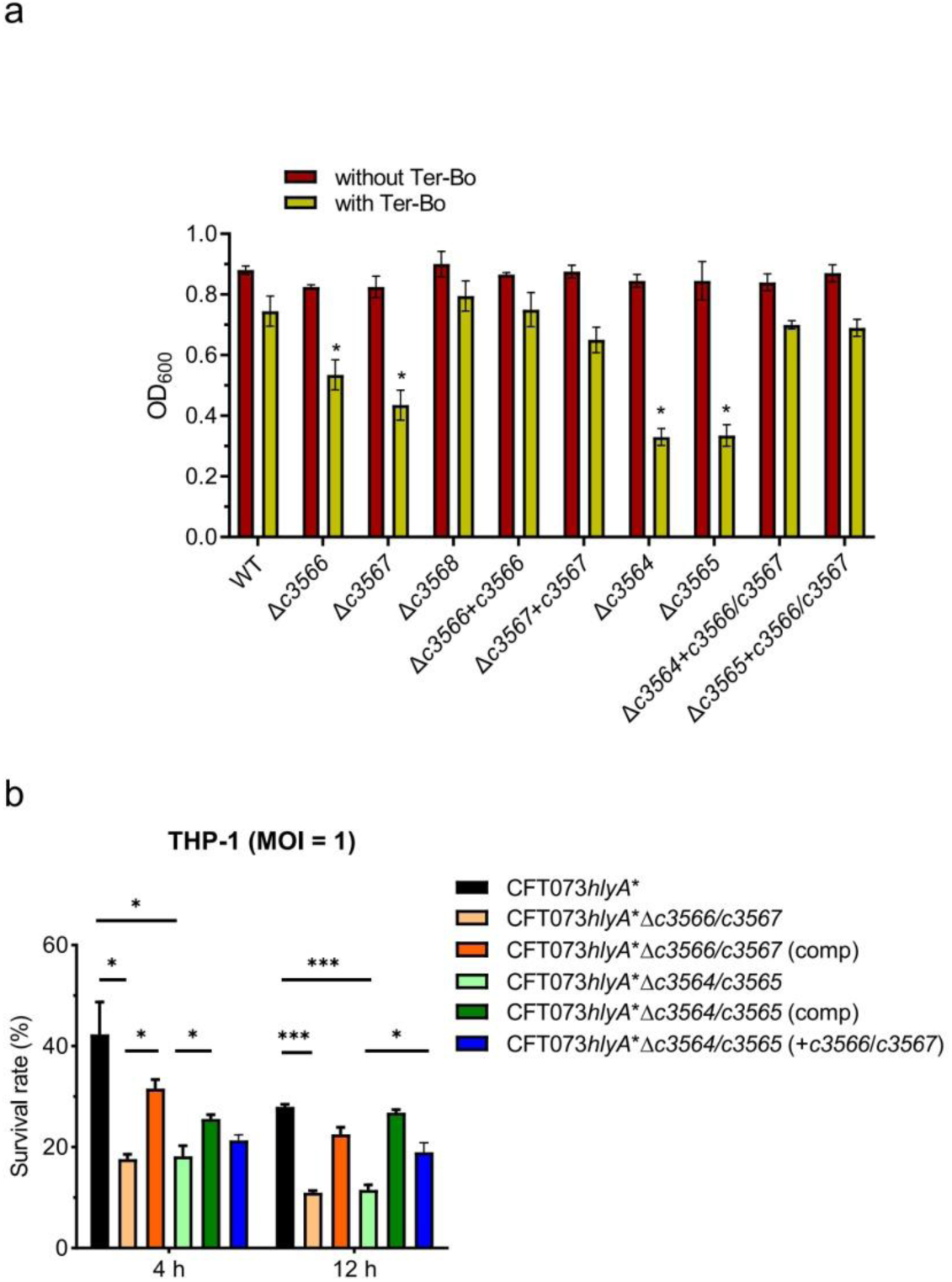
*c3564*/*c3565* and *c3566*-*c3568* mediate oxidative stress resistance and intracellular survival in UPEC. **a**, Oxidative stress resistance assay. Tert-butyl hydroperoxide (Ter-Bo) was added to log-phase bacteria grown in minimal medium at a final concentration of 1 mM, and cultures without added Ter-Bo were used as controls. The OD_600_ values were recorded to assess survival. **b**, Survival of various CFT073 mutant strains within human macrophages. THP-1 cells were infected with human complement-opsonized CFT073 strains at an MOI of 1 for 45 min at 37°C. A gentamicin protection assay was performed to determine the intracellular bacterial counts at the indicated times. Survival was determined as the mean percentage of the number of bacteria recovered at the indicated times compared to that at 1 h after gentamicin treatment, which was considered 100%. CFT073*hlyA*\*, which carries a partially deleted *hlyA* locus, was used as the parental strain, as it exerts very limited cytopathic effects. Comp indicates the strain carries a complementation plasmid. The data represent the mean ± SD from three independent experiments. *, *P* < 0.05; ***, *P* < 0.001 by one-way ANOVA followed by Dunnett’s multiple comparisons test.

Macrophages produce H_2_O_2_ and other oxidants to damage bacterial pathogens^33^. Therefore, we evaluated whether C3564/C3565 and *c3566*-*c3568* are involved in the resistance of UPEC to killing by THP-1 macrophages. As production of HlyA elicits relatively rapid host cell death, thereby preventing long-term incubation of bacteria with host cells (Fig. 3b), we generated a *hlyA* partial deletion strain of CFT073 (Δ amino acids 564-936, herein named CFT073*hlyA*\*) and its variant strains. This partial deletion inactivates hemolysin activities yet allows the monitoring of *hlyA* expression (see the subsection below). Deletion of *c3566*/*c3567* or *c3564*/*c3565* from CFT073*hlyA*\* significantly reduced the intracellular survival of bacteria at 4 and 12 h after the addition of gentamycin (Fig. 5b). Introduction of plasmid-borne *c3566*/*c3567* or *c3564*/*c3565* into the respective mutants rescued intracellular survival, albeit not to wild-type levels. In addition, constitutive expression of *c3566*/*c3567* in the CFT073*hlyA*\*Δ*c3564* or CFT073*hlyA*\*Δ*c3565* mutant strains significantly increased intracellular survival (Fig. 5b). Similarly, *c3566* and *c3567* also contributed to UPEC survival within RAW264.7 murine macrophage cells (Fig. S4). These results indicate that the TCS C3564/C3565 promotes UPEC resistance to killing by macrophages by regulating *c3566* and *c3567* expression.

### C3564/C3565 induces *c3566*-*c3568*-*hlyCABD* expression in response to exogenous H_2_O_2_ and host cell-derived H_2_O_2_

Since C3564/C3565 and its regulated factors C3566/C3567 contribute to UPEC resistance to H_2_O_2_, we investigated whether the expression of *c3566*-*c3568*-*hlyCABD* responds to the presence of H_2_O_2_ in medium. Using a P*_c3566_*-EGFP fusion plasmid for transcriptional analysis, we showed that *c3566*-*c3568-hlyCABD* expression in the wild type was significantly higher in the presence of Ter-Bo than in its absence (Fig. 6a). Deletion of either *c3564* or *c3565* abolished GFP fluorescence, while introduction of *c3564* or *c3565* into the respective mutant restored GFP fluorescence (Fig. 6a), suggesting that *c3564* and *c3565* are required for the induction of *c3566* expression by Ter-Bo.

**Figure 6.**
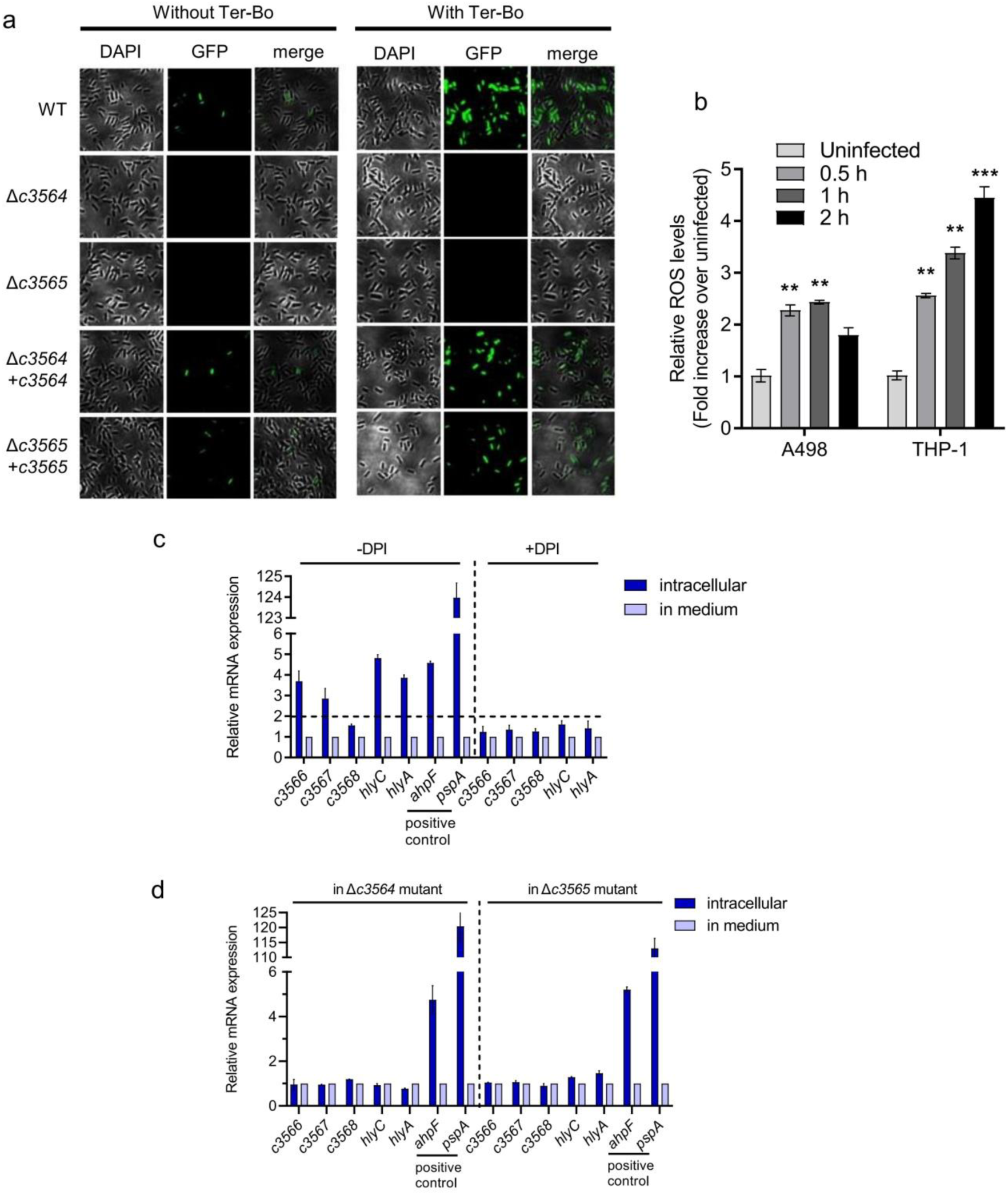
C3564/C3565 induces *c3566*-*c3568*-*hlyCABD* expression in response to *in vitro* and intracellular H_2_O_2_ exposure. **a**, Regulation of *c3566*-*c3568*-*hlyCABD* expression by C3564/C3565 in response to ROS exposure *in vitro*. GFP activity was monitored in wild-type CFT073 and its mutants, all of which carried a P*_c3566_*-EGFP fusion plasmid. Ter-Bo was added at 1 mM. Representative fluorescence micrographs are shown. **b**, ROS levels in A498 and THP-1 cells upon UPEC infection. Cells were infected with wild-type CFT073 for various amounts of time and then treated with H_2_DCFDA (20 μM) for 90 min at 37°C. Cellular fluorescence was assessed using a plate reader (excitation/emission at 485 nm/535 nm), and the levels in the uninfected cells were set to 1. **, *P* < 0.01; ***, *P* < 0.001 by one-way ANOVA followed by Dunnett’s multiple comparisons test. **c**, *c3566*-*c3568*-*hlyCABD* expression induction by intracellular ROS. qPCR was used to examine the transcription levels of *c3566*-*c3568*-*hlyCABD*, and the gene transcription levels in bacteria grown in medium were set as 1. To inhibit the production of ROS, DPI, an inhibitor of NADPH oxidase, was added at 100 μg/mL. The dashed lines represent a fold change cutoff of 2, and a fold change ≥2 was considered significant. **d**, Regulation of *c3566*-*c3568*-*hlyCABD* intracellular expression by C3564/C3565. The transcription levels of *c3566*-*c3568*-*hlyCABD* in the Δ*c3564* and Δ*c3565* mutants within THP-1 macrophages and in medium were tested by qPCR. The gene transcription levels in bacteria grown in medium were set as 1. The positive control genes *ahpF* and *pspA* have been reported to be induced in UPEC inside macrophages.

Then, we hypothesized that the TCS C3564/C3565 responds to intracellular H_2_O_2_ during kidney epithelial cell or macrophage infection. First, we found that ROS production within host cells (A498 and THP-1) was significantly induced at 0.5 h after infection with UPEC. As the infection time increased (up to 2 hpi), ROS levels within THP-1 cells increased, and ROS production was induced to a greater degree in THP-1 cells than in A498 cells (Fig. 6b). We next used CFT073*hlyA*\* to infect THP-1 macrophages for different amounts of time and at different multiplicities of infection (MOIs). The RNA of intracellular bacteria was isolated, and qPCR was used to assess the transcription levels of *c3566*, *c3567, c3568* and *hlyC* and *hlyA*; *ahpF* and *pspA* were used as positive control genes, as the expression of these genes is induced in UPEC within macrophages^34^. As expected, the expression of *ahpF* and *pspA* was dramatically induced in CFT073*hlyA*\* within THP-1 macrophages, and the expression of *c3566*, *c3567*, *c3568*, *hlyC*, and *hlyA* was significantly higher within cells than in medium (Fig. 6c). Diphenyleneiodonium chloride (DPI), an NADPH oxidase inhibitor that represses ROS production, impaired the intracellular expression induction, suggesting that intracellular H_2_O_2_ was involved in the induction of these genes (Fig. 6c). Furthermore, *c3564* deletion and *c3565* deletion each ablated the upregulation of *c3566*, *c3567*, *c3568*, *hlyC*, and *hlyA* within THP-1 cells (Fig. 6d), indicating that within macrophages, *c3564* and *c3565* are essential for H_2_O_2_-mediated induction of the *c3566-c3568-hlyCABD* operon. Additionally, these results were reproduced in the mouse macrophage cell line RAW264.7 under interferon-gamma stimulation (Fig. S5), which results in abundant H_2_O_2_ production^35^. Collectively, these data indicate that within host macrophages, *c3566*-*c3568* and *hlyCABD* expression is linked and is regulated by C3564 and C3565.

## Discussion

We found that the TCS C3564/C3565 directly regulates the expression of *c3566-c3568* and its downstream operon *hlyCABD* because of the cotranscription of *c3566-c3568* and *hlyCABD*. In the presence of H_2_O_2_ (particularly during engulfment by macrophages in which H_2_O_2_ is abundantly produced), *c3566*-*c3568* and hemolysin production is greatly induced. UPEC protect themselves from being killed by H_2_O_2_ in immune cells using a putative Msr system encoded by *c3566-c3567*, serving as a “shield”, and produce large amounts of hemolysin, serving as a “sword”, to trigger rapid host cell death, thus minimizing the detrimental effects that macrophages impose on UPEC and facilitating infection by UPEC. Our study therefore depicts a strategy in which UPEC employ a TCS to coordinate oxidative stress resistance and killing of host cells in response to host-derived H_2_O_2_ signals (Fig. 7). All these functions are encoded in a gene cluster within a pathogenicity island. Our study thus highlights the crucial role of horizontal gene transfer (HGT) in the evolution of bacterial pathogens. In light of their functions, herein we rename *c3564*/*c3565* as *orhK*/*orhR* (<u>o</u>xidative resistance and <u>h</u>emolysis <u>k</u>inase/regulator).

**Figure 7.**
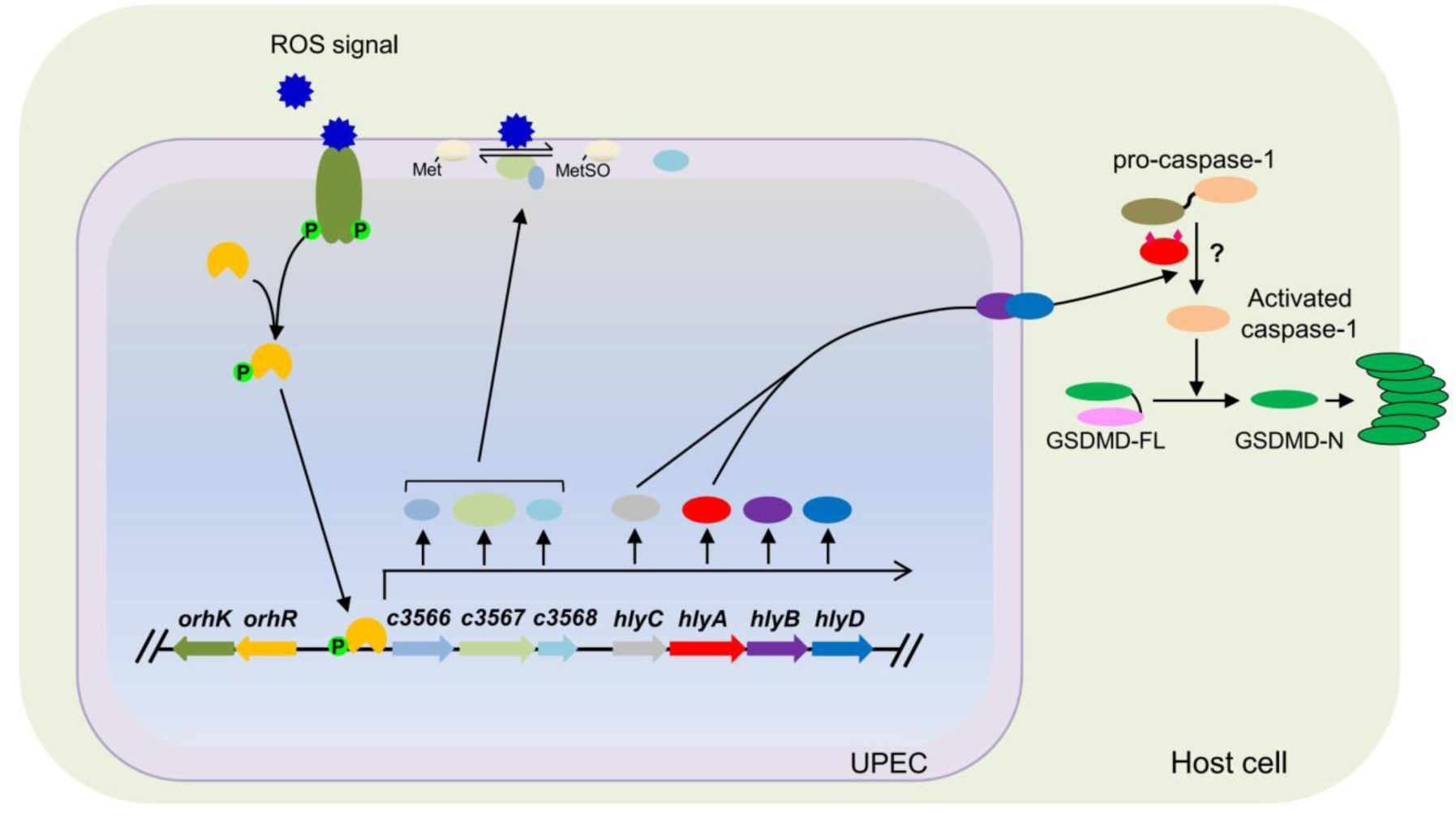
A schematic model of virulence regulation by C3564/C3565. We propose that once inside host macrophages, UPEC is able to sense the presence of ROS through the histidine kinase (HK) C3565, which in turn transphosphorylates the response regulator (RR) C3564; phosphorylated C3564 is an active form and therefore binds to the promoter region of *c3566* and upregulates *c3566*-*c3568*-*hlyCABD* expression. Upon acylation by HlyC, hemolysin encoded by *hlyA* becomes activated, which is translocated by type I secretion machinery containing HlyB and HlyD. C3566 is thought to be membrane-bound, while C3567 and C3568 are periplasmic, and the Msr system C3566-C3567 functions to repair proteins by converting MetSO to Met, thus providing protection. Meanwhile, increased hemolysin production triggers pro-caspase-1 and GSDMD processing, and the resultant GSDMD-N polymerizes and forms pores on the host cell membrane, eventually causing host cell pyroptosis. The question mark (?) indicates a pathway yet to be discovered.

Based on their homology with the YedVW TCS-regulated MsrPQ, C3566 and C3567 encode the membrane-bound and periplasmic components, respectively, of an Msr system. In contrast to MsrPQ, which protects *E. coli* from RCS, the C3566/C3567 system protects UPEC from ROS (Fig. 5). MsrA and MsrB in *E. coli* reduce methionine-*S*-sulfoxide and methionine-*R*-sulfoxide, respectively, and function mainly within the cytoplasm ^36^. The C3566/C3567 system, however, may serve to repair periplasmic proteins and maintain envelope integrity under conditions of oxidative stress, as the substrates of MsrPQ are periplasmic proteins (the chaperone SurA and lipoprotein Pal)^29^. The identities of the C3566/C3567 substrates are currently under investigation in our laboratory. Regarding the roles of *c3566*-*c3568* in UPEC fitness *in vivo*, a prior study showed that inactivation of *c3566*-*c3568*, *c3566* alone or *c3568* alone in CFT073 *dsdA* mutants led to loss of a hypercolonization phenotype, thus establishing an association of *c3566*-*c3568* with UPEC colonization *in vivo*; however, deletion of *c3566* or *c3568* from wild-type CFT073 did not affect UPEC colonization^37^. Therefore, our study identifies a new antioxidative system in UPEC that is potentially involved in colonization *in vivo*.

The role of *c3566-c3568* obviously goes beyond antioxidation; the promoter P*c3566* is necessary for the transcriptional linkage of *c3566-c3568* and *hlyCABD*, and there is presumably no strong terminator after *c3568* to prematurely stop transcription. Both Nhu et al.^22^ and our group (unpublished data) found that insertion of a transposon that carries a CmR gene oriented towards *hlyCABD* within *c3566-c3568* can elevate the expression of *hlyCABD*. These results support our finding that *c3566-c3568* and the *hlyCABD* operon are cotranscribed. Notably, UPEC strains, such as J96 and 536, contain two functional copies of *hlyCABD*, *hly*_I_ and *hly*_II_. A great majority of *hlyA*-positive UPEC isolates in the NCBI database carry the *hly*_I_ copy alone, including CFT073, UTI89, and S65EC^38^. A PCR genotyping analysis of our laboratory collection of 200 UPEC isolates showed that 1) 64 isolates (32%) display hemolytic activities; that 2) 2 isolates (1%) carry two copies and 62 isolates (31%) carry *hly*_I_ only; and that 3) ∼98% of the *hly*_I_-only isolates carry *orhK*/*orhR* and *c3566*-*c3568* operons. These suggest that *orhK*/*orhR*, *c3566*-*c3568* and *hly*_I_ tend to co-occur. Since hemolysin is associated with more severe disease outcomes^39^, the linkage of *c3566*-*c3568* with hemolysin and their regulation by OrhK/OrhR could constitute an important mechanism underpinning severe disease caused by hemolysin-positive UPEC.

It has long been thought that pyroptosis is a form of caspase-1-mediated proinflammatory cell death and that caspase-1 processing sufficiently indicates pyroptosis^40^. Recently, it was found that caspase-11/4/5 can also induce pyroptosis^41, 42^ and that pyroptosis is not necessarily linked to the release of mature IL-1β^43^. Regardless of the caspase activation and cytokine release, increasing lines of evidence show that the real effectors for pyroptosis are the gasdermin-family proteins, primarily GSDMD; upon cleavage, GSDMD-N travels to the membrane and forms pores, allowing the release of cytokines and eventually leading to membrane rupture^44^. UPEC can induce caspase-1/4-dependent inflammatory cell death in bladder epithelial cells^18^. In this study, we further demonstrated that hemolysin induces caspase-1 processing and GSDMD cleavage in both kidney epithelial cells and macrophages, clearly indicating pyroptotic cell death. Hemolysin plays divergent roles in different types of cells from different hosts^39^. This diversity could be at least partially due to different recognition mechanisms of the host cells; some of the various effects exerted by hemolysin are direct, and others are indirect^39^.

Pyroptosis is a defense mechanism by which the host eliminates intracellular pathogens^45^. However, this mechanism can be hijacked by pathogens, and excessive pyroptosis can lead to excessive cytokine levels, cell death, and, eventually, tissue damage/scarring and bacterial dissemination into deeper tissue^45^. Our results show that hemolysin apparently induces pyroptosis in kidney cells and macrophages. Accordingly, reports have indicated that expression of hemolysin is associated with more extensive tissue damage within the urinary tract and more severe clinical outcomes^46^. However, previous work has also elucidated that hemolysin overexpression during acute bladder infection induces rapid and extensive exfoliation and reduces bladder bacterial burdens^47^. Therefore, the balance and timing of hemolysin expression are important in determining disease outcomes. The OrhK/OrhR TCS, together with other reported regulators such as CpxA/CpxR^18^ and FNR^19^, could coordinate to fine-tune hemolysin expression to enhance UPEC fitness during UTI.

In summary, our findings suggest a model in which OrhK/OrhR activates and coordinates the expression of oxidative stress resistance proteins and hemolysin in response to H_2_O_2_ within macrophages (Fig. 7). This regulatory circuit protects UPEC from intracellular oxidative stress and simultaneously promotes hemolysin-mediated pyroptotic cell death of host cells, potentially leading to host tissue damage. Our study provides novel insights into bacterial virulence strategies and suggests OrhK/OrhR as a potential antimicrobial dual target to relieve tissue damage during UTI.

## Methods

### Genetic engineering and construction of recombinant plasmids

Strains and plasmids used in this study are listed in Table S1. DNA amplification, ligation, and electroporation were performed according to standard protocols. All oligonucleotides purchased from Integrated DNA Technologies (Iowa) and Genewiz (Suzhou, China) are listed in Table S2. The various constructs were confirmed by PCR and DNA sequencing (Core Facility, Iowa State University).

Gene deletions in CFT073 were performed using the Lambda Red recombination system described by Datsenko and Wanner^48^. For complementation, the coding regions were amplified from CFT073 genomic DNA and cloned into a modified plasmid (pGEN-P_CmR_) in which the promoter was replaced with the promoter of the chloramphenicol resistance gene from pKD3. For construction of expression plasmids, fragments of the target genes were obtained using the primers in Table S2 and subsequently cloned into pET21a (Novagen) or pGEX-6P-3 (GE Healthcare). P*_c3566_* (from the −35 box to the TSS) was deleted, and the expression of *orhK*/*orhR* was not affected, as determined by qPCR.

### PCR genotyping

The nucleotide sequences of *orhK*, *orhR*, *c3566*, *c3567*, *c3568* and their orthologs in *E. coli* were aligned using the ClustalW2 program. Primers were designed from a conserved region on the basis of G/C content, annealing temperatures, and amplicon sizes. Multiplex PCR was carried out according to the methods of Johnson et al^49^.

### Protein expression and purification

To achieve high expression levels of the target proteins (the cytoplasmic domains of OrhK-GST and His_6_-OrhR), recombinant plasmids were transformed into competent *E. coli* BL21 cells, and their expression was induced by IPTG under optimal growth conditions. The cells were harvested by centrifugation, washed once with lysis buffer, and stored at −80°C until use. The frozen cells were resuspended in lysis buffer containing phenylmethylsulfonyl fluoride and lysozyme and further lysed by sonication. After centrifugation, the supernatant was extracted with a nickel-nitrilotriacetic acid-agarose suspension (Qiagen) or a Pierce GST Spin Purification Kit (Thermo) as described by the manufacturer. Proteins were eluted, recovered and dialyzed against storage buffer^50^. Finally, the protein concentrations were determined with a protein assay kit (Bio-Rad), and the purity was assessed by SDS-PAGE.

### Autophosphorylation and transphosphorylation

Autophosphorylation and transphosphorylation assays were performed as previously described by Yamamoto et al^50^. For autophosphorylation, purified OrhK-GST was diluted with kinase buffer, and the phosphorylation reaction was initiated by adding ATP. The reaction was carried out at 37°C for various amounts of time and terminated by adding an equal volume of SDS-PAGE sample loading buffer. After SDS-PAGE separation, the proteins in the gel were transferred onto a nitrocellulose membrane and detected using a pIMAGO-biotin Phosphoprotein Detection Kit (TYMORA). For transphosphorylation, the phosphorylated form of OrhK prepared as mentioned above was mixed with a mixture of His_6_-OrhR and excess ATP on ice, and the resulting solution was incubated at 37°C for various amounts of time. Reaction termination and subsequent detection of transphosphorylated proteins were performed exactly as mentioned above.

### Cell morphology assay

A498 cells were infected with various bacterial strains in 24-well plates at an MOI of 10 at 37°C, and uninfected cells were used as controls. At 2, 3, and 4 hpi, cell monolayers were visualized with an Axiovert 40C inverted optical microscope (Carl Zeiss), and images were captured with an EOS 1000D Canon camera at 20× magnification.

### LDH cytotoxicity assay

A498, J774A.1, and THP-1 cells were seeded onto 96-well plates until ∼80% confluency was reached. The cells were infected with various bacterial strains at an MOI of ∼10. Cell culture supernatants were collected at 2.5 hpi and subjected to LDH release measurement using an LDH Cytotoxicity Assay Kit (C20301). Cytotoxicity (%) was determined by comparing the LDH in culture supernatants to the total cellular LDH (the amount of LDH released upon cell lysis with 0.1% Triton X-100) according to the formula [(experimental − target spontaneous)/(target maximum − target spontaneous)] × 100^51, 52^.

### Annexin-V/PI staining and flow cytometry analysis

This experiment was carried out as previously described^27^. Briefly, cells were trypsinized, washed with cold PBS, and then resuspended in the binding buffer [10 mM HEPES (pH 7.4), 140 mM NaCl and 2.5 mM CaCl_2_]. One hundred microliter of the cell suspension were transferred into a new tube and stained with 5 μL FITC-Annexin V and 10 μL propidium iodine for 15 min in dark at room temperature. After incubation, 400 μL of binding buffer was added to the solution and further analyzed by FACScallibur flow cytometer (BD Biosciences).

### Blood agar plate assay

Bacterial strains were streaked on sheep blood agar plates (BD Biosciences, NJ), and the plates were incubated at 37°C for 12 h. Colonies of similar sizes were photographed digitally with a stereomicroscope (Olympus) under the same lighting and magnification conditions^18^.

### Transcriptomics by RNA sequencing (RNA-seq)

RNA-seq analysis was performed using a standard protocol, with minor modifications^53^. Briefly, RNA extraction was carried out using a TRIzol (Thermo) method and was followed by determination of the RNA quality and concentration with an Agilent 2100 Bioanalyzer (Agilent Technologies) and a NanoDrop instrument (Thermo Fisher Scientific Inc.), respectively. One microgram of high-quality RNA (A260/A280 ratio ˃ 2.0 and RNA integrity number (RIN) ˃ 7.0) was used for construction of each NextGen sequencing library. Ribosomal RNA was removed from total RNA using a Ribo-Zero rRNA Removal Kit for Bacteria (Illumina). The libraries were constructed according to the manufacturer’s protocol (NEBNext Ultra Directional RNA Library Prep Kit for Illumina). Sequencing of the libraries was performed using a 2×150 paired-end (PE) configuration on an Illumina HiSeq platform according to the manufacturer’s instructions (Illumina, CA); image analysis and base calling were conducted by HiSeq Control Software (HCS) + Off-Line Basecaller (OLB) + GAPipeline-1.6 (Illumina) on the HiSeq instrument. The sequences were processed and analyzed, and the raw reads were assessed with FastQC and further processed with Cutadapt (version 1.9.1). The clean reads were then aligned to the UPEC CFT073 genome (GenBank accession: NC_004431.1) using Bowtie 2 (version 2.1.9, standard options). The reads were counted using HTSeq (version 0.6.1). Differential gene expression analysis was then performed using DESeq2 (version 1.6.3) with R version 3.3.2 following a standard workflow ^53^. All genes with a |log_2_(fold change)|>1 and a Benjamini-Hochberg adjusted *P* value (*q* value) < 0.05 were considered differentially expressed.

### RT-PCR and qPCR

UPEC CFT073 and its derivatives were treated with RNAprotect Bacterial Reagent (Qiagen, CA), and RNA was extracted using an RNeasy Mini Kit (Qiagen) with one-hour in-tube DNase digestion (Qiagen) to remove possible DNA contamination according to the manufacturer’s instructions. One microgram of total RNA was reverse transcribed in triplicate using random hexamers and SuperScript® IV reverse transcriptase (Invitrogen). For the cotranscription test by RT-PCR, primer pairs were designed to span adjacent genes. RNA that was not reverse transcribed served as a negative control, while genomic DNA served as a positive control^25^.

For qPCR analysis, cDNA was used as a template for SYBR Green-based qPCR using TB Green Premix Ex Taq™ II Reagent (Clontech) and an ABI Quant 5 thermocycler (Applied Biosystems). Melting curve analyses were performed after each reaction to ensure amplification specificity. Fold changes in transcript levels were calculated using the 2^−ΔΔCt^ method^54^, and the levels were normalized according to *rpoB* expression. Differences between groups were analyzed by Student’s *t* test. The primers that were used are listed in Table S4.

### EMSAs

To study the binding of OrhR to DNA probes, EMSAs were performed using a commercially available EMSA kit (Invitrogen, CA)^25^. The OrhR-His_6_ fusion protein was purified to homogeneity using Ni-NTA Spin Columns (Qiagen) and dialyzed against binding buffer. The DNA probes were PCR amplified using specific primers and gel purified using a Qiagen MinElute Gel Extraction Kit (Qiagen). EMSAs were performed by adding increasing amounts of purified OrhR-His_6_ fusion protein to a DNA probe (40 ng) in binding buffer (10 mM Tris (pH 7.5), 1 mM EDTA, 1 mM dithiothreitol, 50 mM KCl, 50 mM MgCl_2_, 1 μg/mL bovine serum albumin (NEB)) for a 30-min incubation at room temperature. The reaction mixtures were then subjected to electrophoresis on a 6% polyacrylamide gel in 0.5× TBE buffer (44.5 mM Tris, 44.5 mM boric acid, 1 mM EDTA, pH 8.0) at 200 V for 45 min. The gel was stained for 30 min in 0.5× TBE buffer containing 1× SYBR Gold nucleic acid staining solution (Life Technologies, Grand Island, NY), and then, images were recorded.

### Mapping of TSSs by 5’-RACE

Total RNA was isolated from UPEC strains as described above. TSS mapping was performed using an ExactSTAR Eukaryotic mRNA 5’- and 3’-RACE Kit (Epicentre Biotechnologies) as described in the manufacturer’s guidelines, except that the first two steps prior to the treatment of RNA with tobacco acid pyrophosphatase (TAP) were omitted^55^. The sequences for the primers used can be found in Table S1.

### Growth inhibition by H_2_O_2_ (Ter-Bo)

Overnight cultures of CFT073 and its mutants were diluted 1:100 in M9 minimal medium (containing 0.4% glycerol) and allowed to grow until they reached an optical density at 600 nm (OD_600_) of ∼0.3. To test bacterial resistance to oxidative stress, Ter-Bo was added to the cultures at a final concentration of 1 mM, and cultures without Ter-Bo were used as controls. After incubation for 3 h at 37°C, the H_2_O_2_ resistance was measured by determining the OD_600_^56^.

### Bacterial intracellular survival assay

Macrophages were cultured in 24-well plates and treated with 100 nM phorbol myristate acetate (PMA) for 48 h for differentiation. The cells were infected with human complement-opsonized CFT073 strains at an MOI of 1 for 45 min at 37°C. After infection, the cells were washed three times with PBS and incubated in medium containing 100 μg/mL gentamicin for 1 h to kill extracellular bacteria. The medium was then replaced with maintenance medium containing 10 μg/mL gentamicin for the indicated amounts of time. Afterward, the monolayers were washed with PBS and lysed with 1% Triton X-100, and the released intracellular bacteria were serially diluted and plated on LB agar for enumeration. Survival was determined as the mean percentage of the number of bacteria recovered at the indicated times compared to that at 1 h after gentamicin treatment, which was considered 100%^57, 58^.

### RNA isolation from intracellular UPEC

Bacterial infection and RNA isolation from intracellular bacteria were conducted according to previously reported methods^59^. THP-1 cells (∼10^6^ cells) were seeded in 6-well plates, stimulated with PMA, and then challenged with human complement-opsonized CFT073 strains at an MOI of 10 for 45 min at 37°C. In parallel, bacteria were incubated in cell-free 6-well plates as the control group. After infection, the cells were washed three times with PBS and incubated in medium containing 100 μg/mL gentamicin (DPI was added at this step when needed) for 1 h to kill the extracellular bacteria. The infected macrophages were then lysed in ice-cold RNA stabilization solution (0.2% SDS, 19% ethanol, 1% acidic phenol in water) and incubated on ice for 30 min to prevent RNA degradation. The lysates containing intracellular UPEC were collected and centrifuged, and RNA was extracted from the bacterial pellets with TRIzol (Thermo).

### Immunoblotting

Cells were collected and lysed in 5× SDS loading buffer to obtain protein samples. A standard SDS-PAGE protocol was used to separate the proteins. The separated proteins were transferred to PVDF membranes and detected with an anti-GSDMD or anti-Caspase-1 rabbit monoclonal primary antibody and a horseradish peroxidase-conjugated secondary antibody as previously described^60^. GAPDH was used as a loading control. The band intensities were quantified using ImageJ densitometry analysis.

### ROS measurement

ROS production in macrophages and kidney epithelial cells was measured as previously described (Kaihami et al., 2014), with minor modifications. In 6-well plates, cells were seeded and infected with wild-type CFT073 for the indicated amounts of time. Then, the cells were washed two times with medium and subjected to treatment with 2’,7’-dichlorodihydrofluorescein diacetate (H_2_DCFDA, 20 μM) for 90 min at 37°C. The cells were collected and resuspended in PBS, and the resultant cell suspensions were transferred into 96-well black plates for fluorescence measurement in a plate reader (SpectraMax M3) at an excitation wavelength of 485 nm and an emission wavelength of 535 nm. The mean intensities of uninfected cells were used as controls. The fold changes in fluorescence were calculated by dividing the mean intensity values of the infected groups by those of the uninfected controls.

### Immunostaining and confocal microscopy

Cells (A498 or RAW264.7) were cultured on cell culture dishes, infected with bacterial strains for different time periods, washed, fixed for 15 min with 4% paraformaldehyde in PBS, permeabilized for 20 min in 0.1% Triton X-100 in PBS and blocked using 5% BSA for 1 h. Then, the cells were stained with the indicated corresponding primary antibodies and incubated with fluorescence-conjugated goat anti-mouse IgG (Invitrogen). The nuclei were counterstained with DAPI (Cell Signaling). The slides were mounted using Fluorescence Antifade Mountant (Molecular Probes). Images were captured at room temperature using a confocal microscope (Olympus FluoView FV1000 Confocal System) with a 63× oil immersion objective and Olympus FluoView software (Olympus). The confocal images shown are representative of three independent experiments.

### Statistical analysis

One-way ANOVA (followed by Dunnett’s multiple comparisons test) was used to analyze differences between various mutants and the wild-type strain (GraphPad 9.0, Prism). Student’s *t* test was used for all other binary comparisons. A *P* value < 0.05 was considered to indicate statistical significance.

### Data availability

The data that support the findings of this study are openly available in National Microbiology Data Center (NMDC40002004 to NMDC40002005).

## Acknowledgements

We thank Ashraf H. Hussein for his assistance in mutant construction. This work was partially supported by Natural Science Foundation of Jiangsu Province (BK20191122), the project of Jiangsu Commission of Health (H2019085), and National Natural Science Foundation of China Young Scholars Project (31902242). The funders played no roles in study design, data collection and interpretation, or submission for publication.

## Author contributions

L.G., G.H., C.X., and C.W. conceived and designed the experiments; G.H., C.X., C.W., L.J., and Z.X.s performed the experiments; L.G., G.H., C.X., and C.W. analyzed the data; G.H., C.X., C.W., and L.G. wrote the paper.

## Materials & Correspondence

Correspondences should be addressed to Ganwu Li or Wentong Cai.

