## Supplemental dataset for "A bacterial “shield and sword”: A previously uncharacterized two-component system protects uropathogenic *Escherichia coli* from host-derived oxidative insults and promotes hemolysin-mediated host cell pyroptosis"

### Supplementary materials

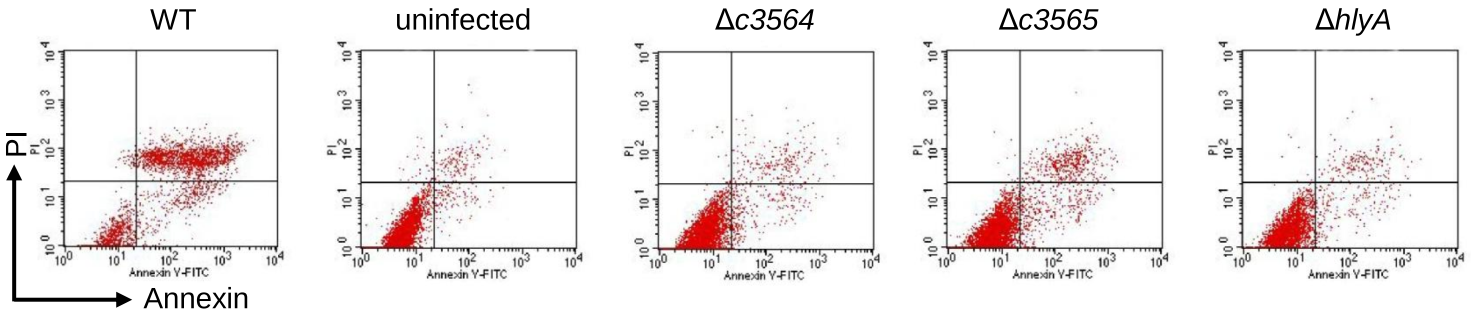

Figure S1. Analysis of the cell death pathways by Annexin-V/PI staining. A498 cells were infected with various CFT073 strains for 1 h, and then stained with Annexin-V and PI, followed by flow cytometry analysis. Live cells are negative for both stains; early apoptotic cells display high Annexin-V signal in the absence of PI staining; necrotic, pyroptotic and late-apoptotic cells have a high PI signal. Shown are typical images of at least two independent experiments.

a

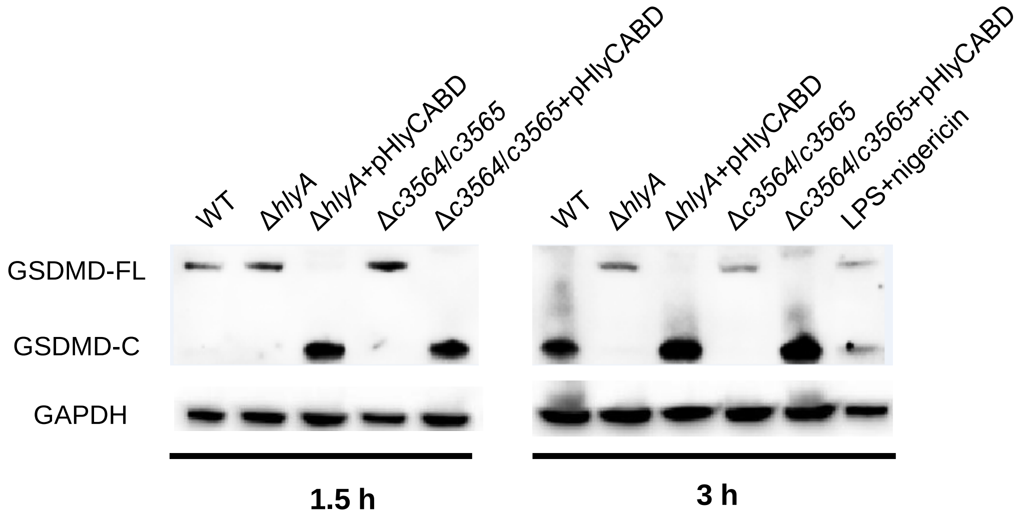

**b**

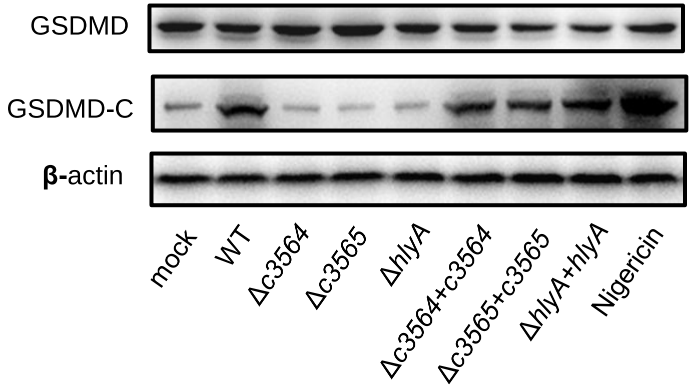

Figure S2. *c3564* and *c3565* promote hemolysin-mediated pyroptosis in human THP-1 and murine J774A.1 macrophages. **a**, *c3564*/*c3565* promotion of hemolysin-mediated processing of GSDMD during infection of murine J774A.1 macrophages**.** LPS plus nigericin, a positive control, were used to induce pyroptosis. **b,** *c3564* and *c3565* promote hemolysin-mediated processing of GSDMD during infection of human THP-1 macrophages. Nigericin, a positive control, was used to induce pyroptosis. All blots are representative of ≥2 independent experiments.

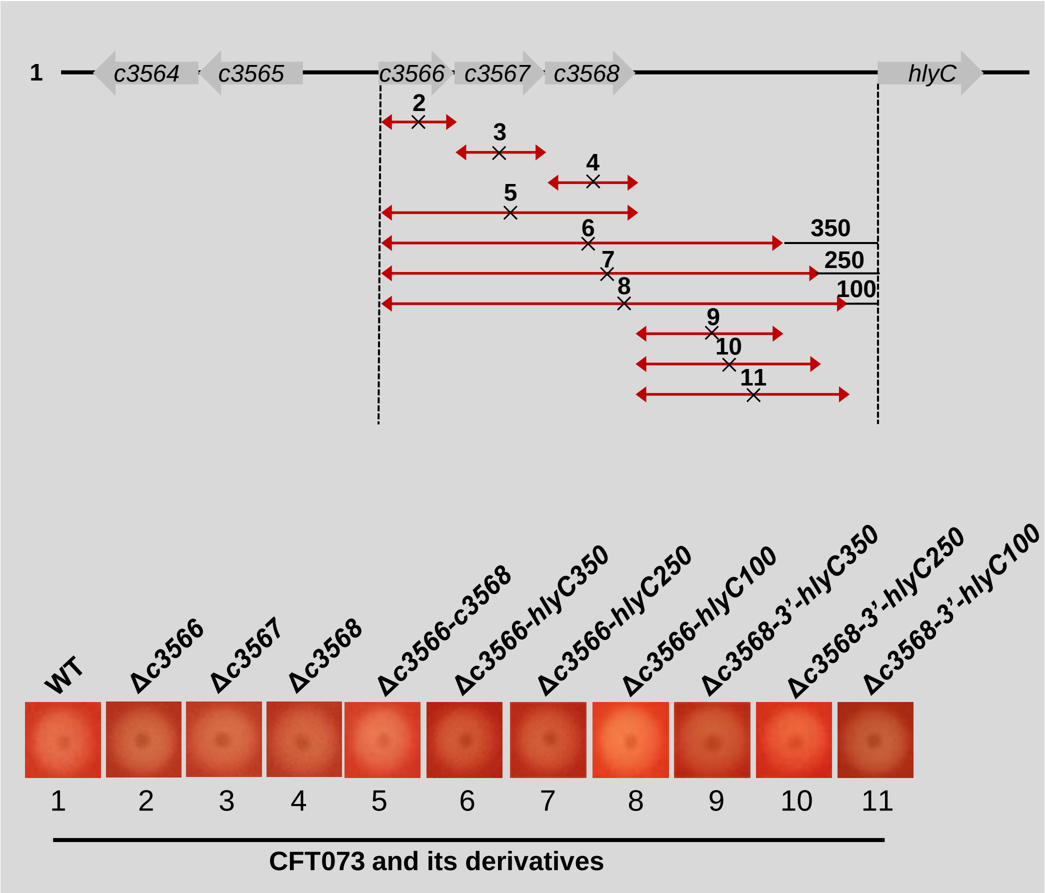

Figure S3. Hemolysis induced by various CFT073 strains on sheep blood agar. Wild-type CFT073 and its deletion mutants are numbered, and the corresponding deleted regions are represented by arrows with crosses in the middle. The dark dot in the center of each image represents a bacterial colony, and the halo around the dark dot represents hemolysis. A wider halo indicates stronger hemolysis. The numbers, 350, 250, and 100, denote the distances between the endpoint of that deletion and the *hlyC* start codon.

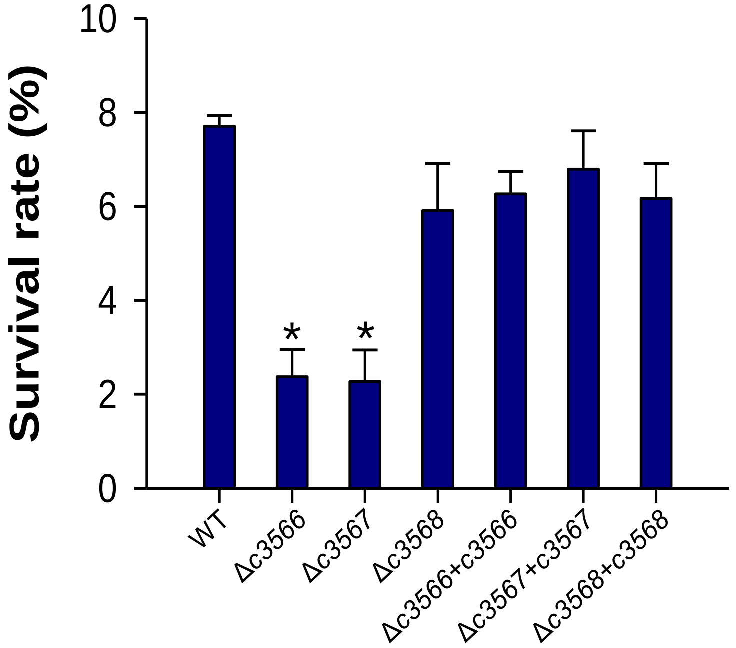

Figure S4. C3566 and C3567 mediate survival of CFT073 within murine RAW264.7 macrophages. RAW264.7 cells were infected with CFT073 strains at an MOI of 1 for 45 min at 37° C. A gentamicin protection assay was performed to determine the intracellular bacterial counts at the indicated times. Survival was determined as the mean percentage of the number of bacteria recovered at the indicated times compared to that at 1 h after gentamicin treatment, which was considered 100%. The data represent the mean ± SD of three independent experiments. *, *P* < 0.05 by one-way ANOVA followed by Dunnett's multiple comparisons test.

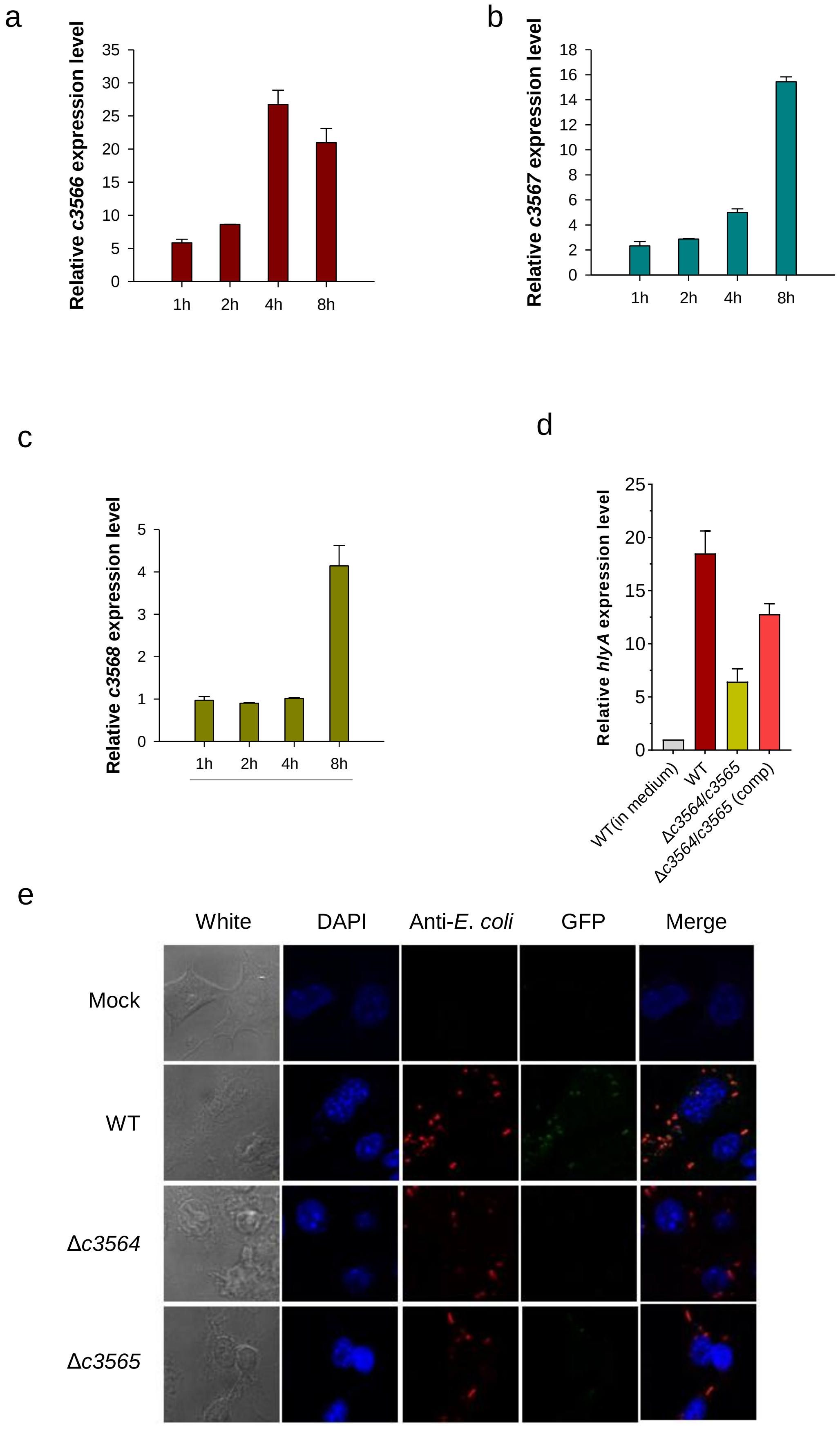
a
Relative c3566 expression level
35
30
25
20
15
10
5
0
1h
2h
4h
8h
b
Relative c3567 expression level
18
16
14
12
10
8
6
4
2
0
1h
2h
4h
8h
c
Relative c3568 expression level
5
4
3
2
1
0
1h
2h
4h
8h
d
Relative hlyA expression level
25
20
15
10
5
0
WT(in medium)
WT
Δc3564/c3565
Δc3564/c3565 (comp)
e
White
DAPI
Anti-E. coli
GFP
Merge
Mock
WT
Δc3564
Δc3565

Figure S5. *c3566-c3568-hlyA* expression is regulated by *c3564* and *c3565* during UPEC intracellular infection of murine RAW264.7 macrophages (primed with interferon-gamma). **a, b, c, and d**, qPCR was used to examine the transcription levels of *c3566* (a), *c3567* (b), *c3568* (c), and *hlyA* (d), and the gene transcription levels in bacteria grown in medium were set as 1. WT, wild-type CFT073. **e**, Regulation of the *c3566* operon by C3564/C3565 within murine RAW264.7 macrophages. GFP activity was monitored in wild-type CFT073 and its mutants, all of which carried a $P_{c3566}$-EGFP fusion plasmid, during infection of RAW264.7 cells. Representative confocal fluorescence micrographs are shown.

**Table S1.** Strains and plasmids used in this study.

| Bacterial strains and plasmids | Genotype/relevant characteristics | Source or Reference |
|---|---|---|
| **Bacterial strains** | | |
| *E. coli* DH5α | Plasmid propagation strain | Invitrogen |
| *E. coli* S17-λpir | RK2 *tra* regulon, *pir*, host for *pir*-dependent plasmids | [1] |
| UPEC CFT073 | Blood isolate from a patient with acute pyelonephritis | [2] |
| Δ*hlyA* | CFT073Δ*lacZYA*Δ*hlyA* | This study |
| Δ*c3564* | CFT073Δ*lacZYA*Δ*c3564* | This study |
| Δ*c3565* | CFT073Δ*lacZYA*Δ*c3565* | This study |
| Δ*c3564*/c*3565* | CFT073Δ*lacZYA*Δ*c3564–65* | This study |
| Δ*c3564*-c*3568* | CFT073Δ*lacZYA*Δ*c3564–68* | This study |
| Δ*c3564–hlyC-350* | CFT073Δ*lacZYA*Δ*c3564–hlyC-350* | This study |
| Δ*c3564–hlyC-250* | CFT073Δ*lacZYA*Δ*c3564–hlyC-250* | This study |
| Δ*c3564–hlyC-100* | CFT073Δ*lacZYA*Δ*c3564–hlyC-100* | This study |
| Δ*c3566*-c*3568* | CFT073Δ*lacZYA*Δ*c3566–68* | This study |
| Δ*c3566–hlyC-350* | CFT073Δ*lacZYA*Δ*c3566–hlyC-350* | This study |
| Δ*c3566–hlyC-250* | CFT073Δ*lacZYA*Δ*c3566–hlyC-250* | This study |
| Δ*c3566–hlyC-100* | CFT073Δ*lacZYA*Δ*c3566–hlyC-100* | This study |
| Δ*c3568–hlyC-350* | CFT073Δ*lacZYA*Δ*c3568–hlyC-350* | This study |
| Δ*c3568–hlyC-250* | CFT073Δ*lacZYA*Δ*c3568–hlyC-250* | This study |
| Δ*c3568–hlyC-100* | CFT073Δ*lacZYA*Δ*c3568–hlyC-100* | This study |
| Δ*c3566* | CFT073Δ*lacZYA*Δ*c3566* | This study |
| Δ*c3567* | CFT073Δ*lacZYA*Δ*c3567* | This study |
| Δ*c3568* | CFT073Δ*lacZYA*Δ*c3568* | This study |
| Δ*c3566*/c*3567* | CFT073Δ*lacZYA*Δ*c3566–67* | This study |
| Δ*c3567*/c*3568* | CFT073Δ*lacZYA*Δ*c3567–68* | This study |
| Δ*P3566* | CFT073Δ*lacZYA*Δ*P3566* | This study |
| Δ*c3564–P3566*::*Pcm* | CFT073Δ*lacZYA*Δ*c3564–P3566*::*Pcm* | This study |
| Δ*c3564–PhlyC*::*Pcm* | CFT073Δ*lacZYA*Δ*c3564–PhlyC*::*Pcm* | This study |
| CFT*hlyA** | *hlyA* partial deletion mutant strain of CFT073 (Δ amino acids 564-936) | This study |
| **Plasmids** | | |
| pMAL-c2X | expression vector | New England Biolabs |
| pET21-a | expression vector | Novagen |

| | | |
|---|---|---|
| pGEX-6P-3 | expression vector | GE Healthcare |
| pGEX-6P-3-c3564 | pGEX-6P-3 carrying partial *c3564* | This study |
| pGEX-6P-3-Mutc3564 | pGEX-6P-3 carrying partial C3564H278A | This study |
| pMAL-*c3564*/c35*65* | pMAL-c2x carrying *c3564–65* under the control of Ptac | This study |
| pMAL-*c3566*-c35*68* | pMAL-c2x carrying *c3566–68* under the control of Ptac | This study |
| pMAL-*c3564* | pMAL-c2x carrying *c3564* under the control of Ptac | This study |
| pMAL-*c3565* | pMAL-c2x carrying *c3565* under the control of Ptac | This study |
| pET21-*c3565* | pET21-a expression vector containing *c3565* gene | This study |
| pET21-*hlyA* | pET21-a expression vector containing *hlyA* gene | This study |
| pGEN-MCS | low copy plasmid for complementation | [5] |
| pHlyCABD | The entire *hlyCABD* cluster plus 1 kb upstream carryied in pGEN-MCS | This study |
| pGEN- $P_{CmR}$ | pGEN-MCS carrying the promoter region of *cat* gene from pKD3 ($P_{CmR}$) | This study |
| p*c3564*/c35*65* | pGEN-MCS carrying *c3564–65* coding region under the control of the native promoter | This study |
| p*c3564* | pGEN-$P_{CmR}$ carrying *c3564* coding region under the control of $P_{CmR}$ | |
| p*c3565* | pGEN-MCS carrying *c3565* coding region under the control of the native promoter | |
| p*c3566* | pGEN-MCS carrying *c3566* coding region under the control of the native promoter | |
| pc35*67* | pGEN-$P_{CmR}$ carrying *c3567* coding region under the control of $P_{CmR}$ | |
| p*c3566*–c35*67* | pGEN-MCS carrying *c3566–67* coding region under the control of the native promoter | This study |
| p*c3564–hlyC* | pGEN-MCS carrying *c3564–hlyC* coding region | This study |
| pEGFP-Prom/hlyC | pEGFP plasmid with promoter replaced with the promoter of *hlyC* | This study |
| pEGFP-Prom/ NC | pEGFP plasmid with promoter replaced with NC sequence | This study |
| pEGFP-Prom/c3566 | pEGFP plasmid with promoter replaced with the promoter of *c3566* | This study |
| pKD3 | template for λ-Red $Chl^r$ cassette | [6] |
| pKD4 | template for λ-Red $Kan^r$ cassette | [6] |

| | | |
|---|---|---|
| pCP20 | encodes FLP recombinase for removal of resistance cassette | [6] |
| pKD46 | λ-Red recombinase expression | [6] |

**Table S2.** Oligonucleotides used in this study.

| Primers | Sequence (5'-3') |
|---|---|
| **For Cloning** | |
| pGEX-6P-c3564-P1 | TCCGAATTCCTACTGGGTAACCCGTCCCAT |
| pGEX-6P-c3564-P2 | CAGGTCGACTCATTATAATGGAAGAAAAAAACG |
| pGEX-6P-c3564-mut-P1 | GGAGATATTGGCAATAAATTCAC |
| pGEX-6P-c3564-mut-P2 | TTTATTGCCAATATCTCCGCTGATTTACGGACGCCATTAAC |
| pGEX-6P-c3564-mut-P3 | CTGTAGGTATCTCAGTTCGGTGTAGGTCGTTCGCTC |
| pGEX-6P-c3564-mut-P4 | AACTGAGATACCTACAGCGTG |
| pET-hlyA-F | GGAATTCCATATGCAGAAGCAAGTCTTTGACCCAT |
| pET-hlyA-R | GAACCGCTCGAGCAGCGTATCATTACCTTTATCACC |
| pET-c3565-F | TCGCGGATCCGAATTCATGAATAACGTAAAAAAATACTG |
| pET-c3565-R | GACGGAGCTCTTATCGCATTCCTGCATTGT |
| phlyCABD-F | TGAAGCTTGGTACCGGGATCCGTTGAGAACTTAAAATTACGTTACGATAAA |
| phlyCABD-R | GAAAGGGCAGATTGTGTCGACTTAACGCTCATGTAAACTTTCTGTTACA |
| pMAL-c3566–68-F | ACGCCATATGCAGCCATTACCGTTAAAACA |
| pMAL-c3566–68-R | ACGCGGATCCTGCGCAGAAATGATGCTTACG |
| pGEN-$P_{CmR}$-F | ACGCGAATTCTAGGAACTTCGGCGCGCCTA |
| pGEN-$P_{CmR}$-R | ACGCCATATGTTTAGCTTCCTTAGCTCCTGA |
| c3564-hlyC-F | ACGCGTCGACTTTATCGACCTCACACGAACA |
| c3564-hlyC-R | ACGCGTCGACTATCCCGGGTTAATAAAGCAT |
| pMAL-c3564–65-F | GGATAACATATGATGAATAACGTAAAAAAAATACTG |
| pMAL-c3564–65-R | CGCCTGATGTTAGGATCCAAGAAAAAAACGAAAGCA |
| pEGFP-Prom/hlyC-P1 | CTGCATTAATGAATCGGC |
| pEGFP-Prom/hlyC-P2 | CCGATTCATTAATGCAGATATTTTAGAGTATACTTGCGCACCCG |
| pEGFP-Prom/hlyC-P3 | GCCCTTGCTCACCATTGCGGTGGCAGGTAAAAAAAG |
| pEGFP-Prom/hlyC-P4 | ATGGTGAGCAAGGGCGAG |
| pEGFP-Prom/NC-P1 | CTGCATTAATGAATCGGC |
| pEGFP-Prom/NC -P2 | CCGATTCATTAATGCAGGACTGATCTTTCAACAGAATACTC |
| pEGFP-Prom/NC -P3 | GCCCTTGCTCACCATCGATTCAAAAAACTTGGATAC |
| pEGFP-Prom/NC -P4 | ATGGTGAGCAAGGGCGAG |
| pEGFP-Prom/c3566-P1 | CTGCATTAATGAATCGGC |

| | |
|---|---|
| pEGFP-Prom/c3566-P2 | CCGATTCATTAATGCAGGCAGATAACGGGTCATCTG |
| pEGFP-Prom/c3566-P3 | GCCCTTGCTCACCATTTGTTTTAACGGTAATGGC |
| pEGFP-Prom/c3566-P4 | ATGGTGAGCAAGGGCGAG |
| **For qPCR** | |
| c3564-rP1 | CGCTTTACCTGAGGGATGA |
| c3564-rP2 | ATACCCGACAGCAGACCAC |
| c3565-rP1 | CATATCAACCGTCTTCGTG |
| c3565-rP2 | CATTCCTGCATTGTCAACT |
| c3566-rP1 | TGGCTTGGTGGTGCGTTAC |
| c3566-rP2 | GGTCCATGACAGCGGGAAA |
| c3567-rP1 | ATGATGCTGACTGGCTGTG |
| c3567-rP2 | AAGGGAATGGAGTGGTAATGT |
| c3568-rP1 | TCATGTTCAATGCTCAGGC |
| c3568-rP2 | TTTTCATCCCATCCGTTTT |
| Hly-c-rp1 | ATTGACTGGATTGCTCCTT |
| Hly-c-rp2 | CCTCCGTGAAATTCTGATA |
| Hly-a-rp1 | GTGACTATCTTTGCACCACAA |
| Hly-a-rp2 | CACTGCCTGCCTTTCCTAA |
| Hly-b-rp1 | AAGTCGGATTGATGTTGAG |
| Hly-b-rp2 | AAATTACGGATCTGGTCTA |
| Hly-d-rp1 | GCCTTTCCTTACACCCGATA |
| Hly-d-rp2 | TGCAGTGACAGCCATACCC |
| **For Deletion** | |
| Del-c3564-F | CTTTTACAAAGGCAGATACACATAACCACCCCAAAATATGCCGCCTGATGgtgtaggctggagctgcttcga |
| Del-c3564-R | TCTGGGGAAAAGGGTATAGGTTTTCAGTTGACAATGCAGGAATGCGATAAcatatgaatatcctccttag |
| Del-c3565-F | ACTGTAAATACTAGGCTTAACCTCTGGCTTAATGTCAGGCTACAATTCATgtgtaggctggagctgcttcga |
| Del-c3565-R | CATCGTCAAATGCTGGGGTAAAATTCAGATAAAGAATATGTGGATAACTTcatatgaatatcctccttag |
| Del-c3564/65-F | CTTTTACAAAGGCAGATACACATAACCACCCCAAAATATGCCGCCTGATGgtgtaggctggagctgcttcga |
| Del-c3564/65-R | CATCGTCAAATGCTGGGGTAAAATTCAGATAAAGAATATGTGGATAACTTcatatgaatatcctccttag |
| Del-hlyA-F | AAAAACAAGACAGATTTCAATTTTTCATTAACAGGTTAAGAGATAATTAAgtgtaggctggagctgcttcga |
| Del-hlyA-R | AATCTTATGTGGCACAGCCCAGTAAGATTGCTATTATTTAAATTAATAAAcatatgaatatcctccttag |
| Del-c3564–68-F | CTTTTACAAAGGCAGATACACATAACCACCCCAAAATATGCCGCCTGATGgtgtaggctggagctgcttcga |

| | |
|---|---|
| Del-c3564–68-R | CGACAGAATATGATGTTTTATCGTAACGTAATTT TAAGTTCTCAACTTATcatatgaatatcctccttag |
| Del-c3564–hlyC-350-F | CTTTTACAAAGGCAGATACACATAACCACCCCAA AATATGCCGCCTGATGgtgtaggctggagctgcttcga |
| Del-c3564–hlyC-350-R | CCAATCAGCTGCCGAATGATGACCTGTGAGTTGT CATGTGAACCTCTCTTcatatgaatatcctccttag |
| Del-c3564–hlyC-250-F | CTTTTACAAAGGCAGATACACATAACCACCCCAA AATATGCCGCCTGATGgtgtaggctggagctgcttcga |
| Del-c3564–hlyC-250-R | AAACCATGCTATTGTATTCTCTTCAATATGCACA TTCTAAAGAACTGTTAcatatgaatatcctccttag |
| Del-c3564–hlyC-100-F | CTTTTACAAAGGCAGATACACATAACCACCCCAA AATATGCCGCCTGATGgtgtaggctggagctgcttcga |
| Del-c3564–hlyC-100-R | TGAGAGAACGATATTAATCCTTTAAATATGTTTC TTGCATTTAGTTTCATcatatgaatatcctccttag |
| Del-c3566–hlyC-350-F | TGATGAACATTCCTTTCCATAATATACTTAGTTTA CAGACAGAGATCACAgtgtaggctggagctgcttcga |
| Del-c3566–hlyC-350-R | CCAATCAGCTGCCGAATGATGACCTGTGAGTTGT CATGTGAACCTCTCTTcatatgaatatcctccttag |
| Del-c3566–hlyC-250-F | TGATGAACATTCCTTTCCATAATATACTTAGTTTA CAGACAGAGATCACAgtgtaggctggagctgcttcga |
| Del-c3566–hlyC-250-R | AAACCATGCTATTGTATTCTCTTCAATATGCACA TTCTAAAGAACTGTTAcatatgaatatcctccttag |
| Del-c3566–hlyC-100-F | TGATGAACATTCCTTTCCATAATATACTTAGTTTA CAGACAGAGATCACAgtgtaggctggagctgcttcga |
| Del-c3566–hlyC-100-R | TGAGAGAACGATATTAATCCTTTAAATATGTTTC TTGCATTTAGTTTCATcatatgaatatcctccttag |
| Del-c3568–hlyC-350-F | ATGGGATGAAAAATGACCACATGGGTAAAAAAG ACAATATGTCTCAATAAgtgtaggctggagctgcttcga |
| Del-c3568–hlyC-350-R | CCAATCAGCTGCCGAATGATGACCTGTGAGTTGT CATGTGAACCTCTCTTcatatgaatatcctccttag |
| Del-c3568–hlyC-250-F | ATGGGATGAAAAATGACCACATGGGTAAAAAAG ACAATATGTCTCAATAAgtgtaggctggagctgcttcga |
| Del-c3568–hlyC-250-R | AAACCATGCTATTGTATTCTCTTCAATATGCACA TTCTAAAGAACTGTTAcatatgaatatcctccttag |
| Del-c3568–hlyC-100-F | ATGGGATGAAAAATGACCACATGGGTAAAAAAG ACAATATGTCTCAATAAgtgtaggctggagctgcttcga |
| Del-c3568–hlyC-100-R | TGAGAGAACGATATTAATCCTTTAAATATGTTTC TTGCATTTAGTTTCATcatatgaatatcctccttag |
| Del-c3566-F | ATGAACATTCCTTTCCATAATATACTTAGTTTAC AGACAGAGATCACATAgtgtaggctggagctgcttcga |
| Del-c3566-R | CTGAACGGGTTTCCATAAAACCAGACCAGACAA TAGCAGAGCAGCGCCATcatatgaatatcctccttag |
| Del-c3567-F | CTGACTATACTATTCGGTGGTTATGATACAGTGC GTTATATCCACTTTCTgtgtaggctggagctgcttcga |

| | |
|---|---|
| Del-c3567-R | CCTCCTCATTTTTAACAATTGTATCAACAACCAC CAAACCAGTTATAACCcatatgaatatcctccttag |
| Del-c3568-F | ATAACTGGTTTGGTGGTTGTTGATACAATTGTTA AAAATGAGGAGGAACCgtgtaggctggagctgcttcga |
| Del-c3568-R | AAAACGACAGAATATGATGTTTTATCGTAACGTA ATTTTAAGTTCTCAACcatatgaatatcctccttag |
| Del-c3566/67-F | ATGAACATTCCTTTCCATAATATACTTAGTTTAC AGACAGAGATCACATAgtgtaggctggagctgcttcga |
| Del-c3566/67-R | CCTCCTCATTTTTAACAATTGTATCAACAACCAC CAAACCAGTTATAACCcatatgaatatcctccttag |
| Del-c3567/68-F | CTGACTATACTATTCGGTGGTTATGATACAGTGC GTTATATCCACTTTCTgtgtaggctggagctgcttcga |
| Del-c3567/68-R | AAAACGACAGAATATGATGTTTTATCGTAACGTA ATTTTAAGTTCTCAACcatatgaatatcctccttag |
| Del-c3566–68-F | ATGAACATTCCTTTCCATAATATACTTAGTTTAC AGACAGAGATCACATAgtgtaggctggagctgcttcga |
| Del-c3566–68-R | AAAACGACAGAATATGATGTTTTATCGTAACGTA ATTTTAAGTTCTCAACcatatgaatatcctccttag |
| Del-P3566-F | GCAGATTAAGCCGCAACAGAGCTGCAATATCAT AATCATCTTCCATCAGCgtgtaggctggagctgcttcga |
| Del-P3566-R | ACAGAGGATGTATTTGTTTTAACGGTAATGGCTG CATTATGTGATCTCTGcatatgaatatcctccttag |
| Del-c3564–P3566::Pcm-F1 | CTTTTACAAAGGCAGATACACATAACCACCCCAA AATATGCCGCCTGATGgtgtaggctggagctgcttcga |
| Del-c3564–P3566::Pcm-R1 | tccgtcacaggtaggcgcgccatatgaatatcctccttag |
| Del-c3564–P3566::Pcm-F2 | ctaaggaggatattcatatggcgcgcctacctgtgacgga |
| Del-c3564–P3566::Pcm-R2 | CAACAGCGTAACCACAGAGGATGTATTTGTTTTA ACGGTAATGGCTGCAttttagcttccttagctcctga |
| Del-c3564–PhlyC::Pcm-F1 | CTTTTACAAAGGCAGATACACATAACCACCCCAA AATATGCCGCCTGATGgtgtaggctggagctgcttcga |
| Del-c3564–PhlyC::Pcm-R1 | TCCGTCACAGGTAGGCGCGCCATATGAATATCCT CCTTAG |
| Del-c3564–PhlyC::Pcm-F2 | CTAAGGAGGATATTCATATGGCGCGCCTACCTGT GACGGA |
| Del-c3564–PhlyC::Pcm-R2 | CAGAGCCAGGATACATGCCCAAGAACCTCTAAT GGATTGTTCATATTCAttttagcttccttagctcctga |
| Del-hlyA*-F | TGTTACGCCATTGTTAACTCCCGGTGAGGAAATT CGTGAAAGGAGGCAGTCCGGAgtgtaggctggagctgcttc ga |
| Del-hlyA*-R | TATTATTACTCTGTTGATACTCAAGTGCCTTTTTA AGGGAATCTGGTGTGATTAC catatgaatatcctccttag |

**For Mutation verification**

| | |
|---|---|
| Check-c3564–65-F | CGACCTCACACGAACAACGA |
| Check-c3564–65-R | GCGAAGTGCCACAGTAACGC |
| Check-c3564-F | ACAGGTTATCCCGGTGTC |
| Check-c3564-R | TCAAAGCGTTGTTTCGTC |
| Check-c3565-F | TTTCAGATCATCCCTCAG |
| Check-c3565-R | GCTACAAATTCCCAGAGT |
| Check-c3566-F | TTTACCCCAGCATTTGACG |
| Check-c3566-R | CAGAAGGGAAAACTGAACG |
| Check-c3567-F | GCTCTGCTATTGTCTGGTCT |
| Check-c3567-R | TTTCATGGTTCCTCCTCAT |
| Check-c3568-F | TGGCGACAGCACTACATTC |
| Check-c3568-R | CCGATACAGAGCCTGACATT |
| Check-c3564–68-F | CGACCTCACACGAACAACGA |
| Check-c3564–68-R | TGCGCAGAAATGATGCTTACG |
| Check-c3564–hlyC-F | CGACCTCACACGAACAACGA |
| Check-c3564–hlyC-R | CCCGAAAGGAGCAATCCAGT |
| Check-P3566-F | TTGCCCCATGGCTTCCTGAC |
| Check-P3566-R | GCGAAGTGCCACAGTAACGC |
| Check-c3566–68-F | ATGGAAAAGGGTTTCGGTAGAT |
| Check-c3566–68-R | TGCGCAGAAATGATGCTTACG |
| Check-c3566–hlyC-F | TTGCCCCATGGCTTCCTGAC |
| Check-c3566–hlyC-R | CCCGAAAGGAGCAATCCAGT |
| Check-c3568–hlyC-F | TGAGCAAAACGGATGGGATGA |
| Check-c3568–hlyC-R | CCCGAAAGGAGCAATCCAGT |
| Check-hlyA-F | CTGGGCCAGTTCCCCATTAC |
| Check-hlyA-R | GGGGATTTCGTTGCTCCAGA |
| Check-hlyA*-F | GTGTCACCAGAAATGGAGAC |
| Check-hlyA*-R | ATTAAGATTATCCTGACTTCC |
| **For EMSA** | |
| Prom-c3565-F | CAGCGTAACCACAGAGGATG |
| Prom-c3565-R | CGCAACAGAGCTGCAATATC |
| Control-c3565-F | ATGACGTGCAAACCAGAGCA |
| Control-c3565-R | GTCCTGGGACTGGAAATGGG |
| Prom-c3566-F | TCCCAGGTCTGCTTGTCTAGT |
| Prom-c3566-R | CCTGCGACAGTAACAGCAT |
| Control-c3566–67-F | TCTGCTGATTAATGCCACGA |
| Control-c3566–67-R | CAATGCACGCGGAACCAATA |
| **For 5'-RACE** | |
| Race-RNA-adapter | UCAUACACAUACGAUUUAGGUGACACUAUAGA GCGGCCGCCUGCAGGAAA |
| Race-adapter-F | GCGC<u>GAATTC</u>ACACATACGATTTAGGTGACACT |
| Race-GSP-c3566 | GTCA<u>GTCGAC</u>GTGCCTGCGTAAGGGATTCT |
| MultiP PCR-c3568–hlyC-F | ACGGATGGGATGAAAAATGA |

| | |
|---|---|
| MultiP PCR-c3568–hlyC-R | TGCCCAAGAACCTCTAATGG |
| **For RT-PCR** | |
| RT-c3566–c3567-F | CCCGCTGTCATGGACCTCAC |
| RT-c3566–c3567-R | TCACGTACCAGCCCCCGTAT |
| RT-c3567–c3568-F | CGCCGGACAGACGTCGTTTA |
| RT-c3567–c3568-R | TTGTGTCTGCAGCCTGAGCAT |
| RT-c3568–hlyC-F | TGAGCAAAACGGATGGGATGA |
| RT-c3568–hlyC-R | CCCGAAAGGAGCAATCCAGT |
| RT-hlyC–hlyA-F | CTGGGCCAGTTCCCCATTAC |
| RT-hlyC–hlyA-R | CTCTGCTGTGCCGAATACCT |
| RT-hlyA–hlyB-F | GTTGTACGGCAGTGAGGGAG |
| RT-hlyA–-hlyB-R | AGATTTCGCAGCAAGCAACC |
| RT-hlyB–hlyD-F | CTGGTTACGTCGTCAGGTGG |
| RT-hlyB–hlyD-R | AACGTATCAGCTTCAGCTCCC |
